# Cyanobacterial α-carboxysome carbonic anhydrase is allosterically regulated by the Rubisco substrate RuBP

**DOI:** 10.1101/2023.07.31.551272

**Authors:** Sacha B. Pulsford, Megan A. Outram, Britta Förster, Timothy Rhodes, Simon J. Williams, Murray R. Badger, G. Dean Price, Colin J. Jackson, Benedict M. Long

## Abstract

Cyanobacterial CO_2_ concentrating mechanisms (CCMs) sequester a globally significant proportion of carbon into the biosphere. Proteinaceous microcompartments, called carboxysomes, play a critical role in CCM function, housing two enzymes to enhance CO_2_ fixation: carbonic anhydrase (CA) and Rubisco. Despite its importance, our current understanding of the carboxysomal CAs found in ɑ-cyanobacteria, CsoSCA, remains limited, particularly regarding the regulation of its activity. Here, we present the first structural and biochemical study of CsoSCA from the cyanobacterium *Cyanobium PCC7001*. Our results show that the *Cyanobium* CsoSCA is allosterically activated by the Rubisco substrate ribulose-1,5-bisphosphate (RuBP), and forms a hexameric trimer of dimers. Comprehensive phylogenetic and mutational analyses are consistent with this regulation appearing exclusively in cyanobacterial ɑ-carboxysome CAs. These findings clarify the biologically relevant oligomeric state of α-carboxysomal CAs and advance our understanding of the regulation of photosynthesis in this globally dominant lineage.

**One-Sentence Summary:** The carboxysomal carbonic anhydrase, CsoSCA, is allosterically activated by the Rubisco substrate RuBP, revealing a novel mechanism controlling key enzyme activity in cyanobacterial α-carboxysomes.

## INTRODUCTION

A myriad of CO_2_ concentrating mechanisms (CCMs) have independently evolved to promote the rapid and efficient reduction of atmospheric CO_2_ into organic compounds. CCMs work to increase the local concentration of CO_2_ near ribulose-1,5-bisphosphate carboxylase/oxygenase (Rubisco), the primary carboxylase of the Calvin–Benson–Bassham (CBB) cycle, thereby increasing its substrate turnover and competitively inhibiting off-target oxygenation reactions^1–4^. These systems are an essential component of the global carbon cycle, catalysing about half of global photosynthesis^2, 5^. The bacterial CCM, found in all cyanobacteria and some autotrophic bacteria, consists of two key elements: first, energy-coupled inorganic carbon (C_i_; primarily HCO_3_^-^ and CO_2_) transporters actively establish a concentrated pool of HCO_3_^-^ within a cytosol lacking free carbonic anhydrases (CAs); this HCO_3_^-^ then diffuses into proteinaceous microcompartments called carboxysomes that house CA and Rubisco^2, 3^. Here, the CA converts HCO_3_^-^ to CO_2_ to elevate luminal CO_2_, promoting Rubisco-catalysed CO_2_ reduction^6^. Bacterial CCMs have arisen in two distinct lineages: β-carboxysomes are found exclusively in β-cyanobacteria, containing Form1B Rubisco with component genes encoded by the *ccm* operon and satellite loci^7^, whereas α-carboxysomes are found in photoautotrophic α-cyanobacteria and several bacterial chemoautotrophs, and are distinguished by the presence of FormIA Rubisco and the clustering of carboxysome-associated genes into a discrete *cso* <u>c</u>arboxy<u>s</u>ome <u>o</u>peron^8^.

α-Cyanobacteria are a globally dominant photoautotrophic lineage across marine and freshwater systems, encompassing two of the most abundant photosynthetic taxa on Earth (*Synechococcus* and *Prochlorococcus*)^5, 9^. Although their global mass is a fraction of that of plant systems, they are estimated to contribute around 25% of global primary production, i.e. carbon fixation^5, 8^. Regulation of carbon fixation is essential for effective energy production. Indeed, the CCM is a striking example of how cells may induce changes in physiological state in response to environmental conditions. This adaptive capacity is a critical feature of these processes, involving regulation at the transcriptional and protein level, allowing the bacterial CCM to competitively support life in a range of ecological contexts^10, 11^. For example, the CCM-related Ci transporter is regulated by gene expression and allosteric effectors^12–14^. Likewise, Rubisco content is transcriptionally regulated and its activity modulated by activases^15–19^. Finally, the carboxysome composition and morphology itself is responsive to environmental cues^20–24^. However, little is known of how, or whether, carbonic anhydrase, the other enzymatic component of the carboxysome, is regulated^25^.

The fundamental role of CAs in photosynthesis is well established^6^. This versatile protein superfamily catalyses the reversible hydration of CO_2_ [CO_2_ + H_2_O ⇌ HCO_3_^-^ + H^+^], comprising eight reportedly evolutionarily distinct classes (α, β, γ, δ, θ, η, ζ and ι), distributed across the tree of life in a kingdom-nonspecific manner^26^. In many cases, the enzyme directly supplies Rubisco with CO_2_, promoting its efficient reduction by ensuring reaction rate optimization through a constant, high concentration of the enzyme-substrate complex^6, 27^. Controlling Rubisco activity through buffering a bicarbonate pool in this way optimizes carbon fixation, coordinating CO_2_ assimilation rates with the generation of NADPH/ATP produced in light reactions^27^. Correspondingly, microbial and biochemical studies have established an absolute requirement for CA activity within the carboxysome^28, 29^. The α-carboxysome contains a highly divergent β-CA known as CsoSCA, characterisation of which has occurred exclusively through the isoform from the chemoautotroph *H. neapolitanus*^30, 31^. Indeed, structural and compositional studies have revealed sequence variation between cyanobacterial and proteobacterial carboxysome components, and distinct carboxysome organisation between these taxa^32–37^. Given the differences in underlying metabolism between these photo- and chemoautotrophs, this has restricted our understanding of the CA-Rubisco feedback in ɑ-cyanobacterial carboxysomes.

Here, we present a detailed biochemical, structural, and evolutionary analysis of a CsoSCA from a photoautotrophic cyanobacterium, *Cyanobium* PCC7001 (*Cyanobium*), revealing previously unknown aspects of this isoform’s activity and molecular structure that form the basis for carboxysome regulation and organisation. We found that, unlike the CA from the chemoautotrophic bacterium *H. neapolitanus* (*Hn*CsoSCA), the *Cyanobium* isoform (*Cy*CsoSCA) is regulated by the Rubisco substrate, ribulose-1,5-bisphosphate (RuBP) for activity, which constitutes a feedback loop in the metabolic pathway to optimize the use of carbon. Detailed evolutionary analysis extends this, revealing the sequence motifs for this regulation are not found in chemoautotrophic bacteria, and expanding our understanding of ɑ-cyanobacterial photosynthesis and CCM diversification more broadly.

## RESULTS

### Cyanobium CsoSCA requires RuBP for activity

Despite homology to the constitutively active *Hn*CsoSCA^30^, in our hands *Cy*CsoSCA did not show detectable HCO_3_^-^ dehydration/CO_2_ hydration activity under standard assay conditions^20^. This surprising result indicated a potential additional requirement for *Cy*CsoSCA function. Given the previous observation of *Cyanobium* carboxysome function *in vitro*^20^, and the established reliance on CA activity^6, 28, 38^, we assessed *Cy*CsoSCA function under Rubisco assay conditions, where RuBP and Mg^2+^ are the key additional components. The addition of RuBP resulted in the concentration-dependent activation of *Cy*CsoSCA, with a *K*_M_ for RuBP of 18 µM, in a similar range as the *Cyanobium* Rubisco *K*_M_ for RuBP (36 µM)^38^. Comparatively, *Hn*CsoSCA activity was unaffected by RuBP. Above 100µM RuBP *Cy*CsoSCA activity rates match those recorded for *Hn*CsoSCA (Fig. 1A). Notably, the *Cy*CsoSCA RuBP response curve is best described by a sigmoidal Hill function (R^2^ = 0.99), typically indicative of an allosteric activation mechanism.

**Figure 1.**
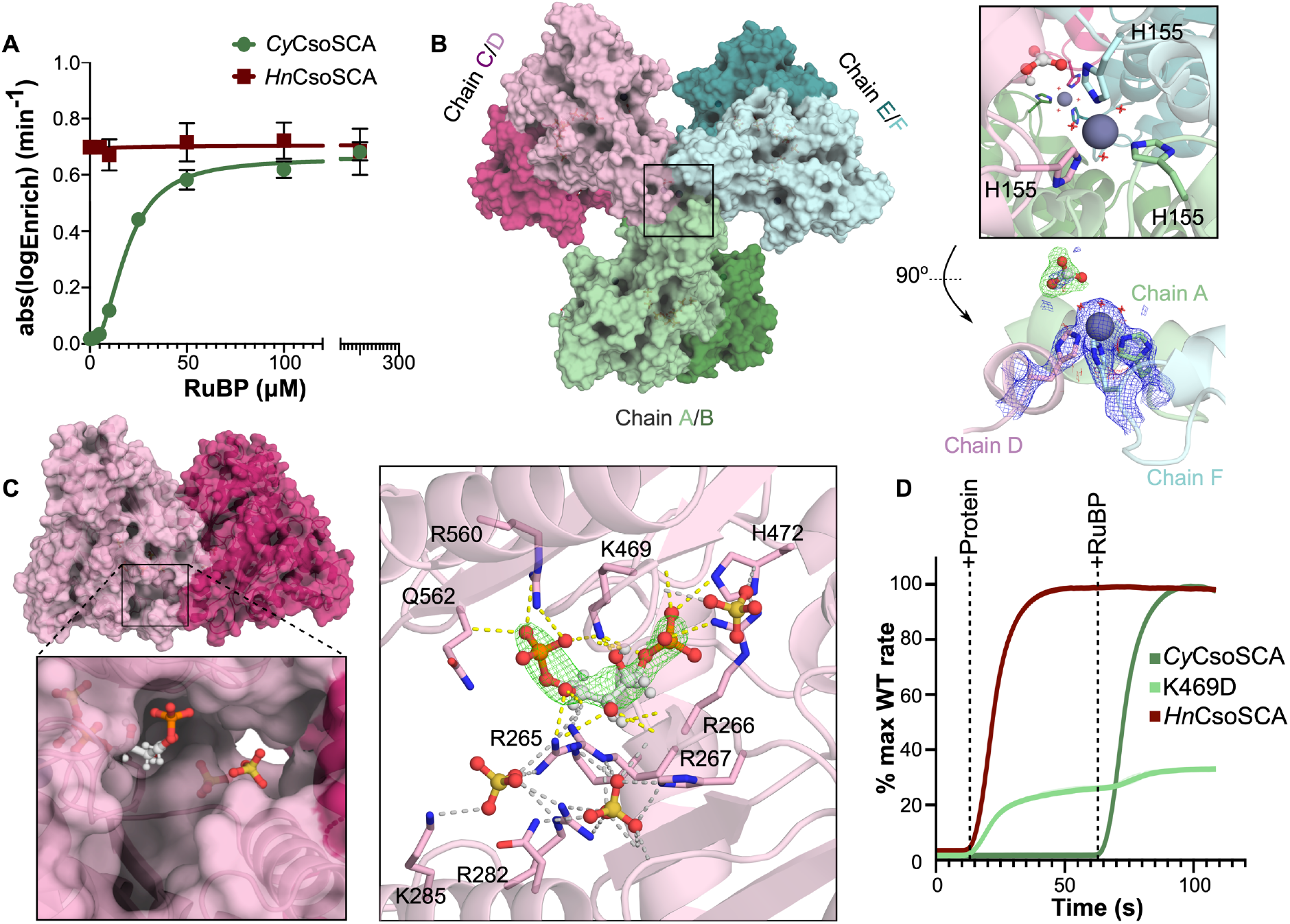
RuBP allosterically regulates CyCsoSCA. **A** CyCsoSCA and HnCsoSCA activity as a function of RuBP concentration, measured using Membrane Inlet Mass Spectrometry (MIMS). Measurements reported are an average of three technical replicates, error bars represent standard error. Curves fitted using GraphPad Prism. **B** The homohexameric CyCsoSCA structure solved to 2.3Å (PBD: 8THM). The ‘trimer of dimers’ arrangement is shown with dimers coloured pink, green or blue, monomers indicated as different shades, chains annotated. The black square denotes the apical structural zinc atoms with further detail shown in the right insert. Waters in the octahedral coordination sphere are shown as red crosses. Zinc ions are shown as grey spheres, an opportune bicarbonate ion is shown in ball and stick representation. Below: 2m*F*_o_-d*F*_c_ density at key interacting residues is shown (1s). Polder omit map (green) is shown at a contour level of 5.0 σ to highlight bicarbonate ion density at the A/D/F apex. **C** The C/D dimer is shown with a box and insert highlighting the RuBP binding pocket of Chain D (light pink monomer). Right panel: The RuBP binding site is shown with Polder omit map density^39^ of the ligand overlayed at a contour level of 7.0 σ. Sulfate ions and RuBP shown in ball and stick representation. Polar interactions are shown as dotted lines, those corresponding to interactions with RuBP directly are coloured yellow, other secondary/SO_4_ interactions shown in grey. **D** The relative rate of CyCsoSCA, the K469D mutant of CyCsoSCA, and HnCsoSCA are shown over time as a proportion of the maximum recorded reaction rate for each respective wild type enzyme. Time points at which protein and 100µM RuBP were added to the MIMS cuvette are indicated. Curve is the mean of three technical replicates with shaded area around the curve representative of standard deviation from the mean (not visible due to scale).

### Cyanobium CsoSCA Structure reveals RuBP binding site and novel oligomeric state

To understand the structural basis for RuBP activation, CsoSCA was co-crystallised with RuBP. Diffracting crystals were obtained almost exclusively in saturating levels of RuBP with the final CsoSCA crystal structure solved through molecular replacement to a resolution of 2.3 Å (Table S1). The resulting structure showed a homohexameric trimer of dimers consistently arranged in the asymmetric unit with P212121 symmetry (Fig. 1B). Size-Exclusion Chromatography (SEC) corroborates both *Cy*CsoSCA and *Hn*CsoSCA are primarily hexameric in solution (Fig. S10), contrasting with previous observations^30^. While the *Cy*CsoSCA dimer interface is highly reminiscent of *Hn*CsoSCA^30^, additional contacts at the N-terminal domain (NTD) of each monomer mediate further quaternary assembly, forming two apices of the final hexamer (Fig. 1B). A metal ion is evident at each apex, coordinated by a His_3_(H_2_O)_3_ octahedral coordination sphere, comprising His155 donated by a distinct monomer (Fig. 1B). This residue sits within a helical bundle in the NTD denoted here as the ‘hook motif’. The electron density and coordination geometry are consistent with a zinc ion. Density corresponding to a HCO_3_^-^ ion was observed at one of the trimer apices. While the pH of crystallisation conditions favours bicarbonate, this species was not present at saturating levels in the crystallisation conditions, which could explain its absence at the opposing apex.

Density consistent with RuBP was observed in all monomers within a positively charged pocket near the dimer interface that extends into the protein core (Fig. 1C, S2 and S8; Table S2 and S3). While variations in omit map density at these positions were observed, given the consistency of this density across each chain, the dependence on RuBP for crystallization, and the observation RuBP was an allosteric activator of CA (Fig. 1), we were confident in the modelling of RuBP at this site. Two sulfate ions, likely from the crystallisation solvent, could be modelled with high confidence at the entrance of this site in all monomers. While slight variations in RuBP ligand conformation are evident in each chain, contacts at Arg266, Lys469 and Arg560 are consistently observed (Fig. S2 and S3). Most notably, RuBP curls around Lys469, mediating multiple H-bonds with the ligand. To confirm that this region is responsible for RuBP binding, we mutated Lys469 to an Asp, the amino acid at the corresponding position in the constitutively active *Hn*CsoSCA isoform. This results in a biphasic activity profile with detectable CA activity evident in the absence of RuBP and a minor increase in activity upon addition of the ligand (Fig. 1D). This directly implicates K469 in the RuBP-mediated activation mechanism and further supports this region as the RuBP binding pocket.

### RuBP regulation of Cyanobium CsoSCA is allosteric

Given the sigmoidal *Cy*CsoSCA activation curve and RuBP binding site distinct from the active site (Fig.1), we hypothesised RuBP acts as an allosteric activator. The β-CA family is the only CA family known to exhibit allostery to date^40^. Alignments between *Cy*CsoSCA and a structure of a previously characterised Type II β-CA bound to the allosteric bicarbonate ion^40^ show RuBP engages distinct residues and sits further from the active site, indicating a distinct regulatory mechanism (Fig. S4). The RuBP site overlays with the region of the CsoSCA C-terminal domain (CTD) that has lost the second symmetric catalytic zinc site seen in canonical β-CAs, likely following gene duplication and divergence of the catalytic domain^30^. A structural analysis of the *Cy*- and *Hn*CsoSCAs was conducted to identify potential allosteric networks within the cyanobacterial variant. *Cy*CsoSCA monomers align well with the canonical *Hn*CsoSCA (37.2% sequence identity, Cα RMSD of 1.5 Å, Fig. 2A) and the three domains (NTD, Catalytic domain and CTD)^30^ are evident (Fig. 2A). The Cys_2_His(H_2_O) tetrahedral coordination of the catalytic zinc ion typical of β-CAs is maintained and the overarching active site is highly homologous between each CsoSCA isoform. The Asp-Arg dyad between active site residues Asp246 and Arg248 precludes the inactive Cys_2_HisAsp coordination sphere across all *Cy*CsoSCA monomers, reinforcing the classification of CsoSCA as Type I^30^. This type of β-CA has not previously been associated with allostery. Manual inspection of H-bonds within the protein identified a network linking the catalytic Asp246 backbone and the Arg266 sidechain that in turn binds RuBP, mediated by a water molecule and Leu249 backbone groups (Fig. 2B). In the corresponding region in the constitutively active *Hn*CsoSCA, Lys179 (Ala250 in *Cy*CsoSCA) occupies this space, coordinating a more extensive interaction network reinforced by multiple water molecules. Notably, Arg196 (Arg266 in *Cy*CsoSCA) binds Asp409 (Lys469 in *Cy*CsoSCA) identified above as a key determinant in RuBP-dependent activity (Fig. 2C).

**Figure 2.**
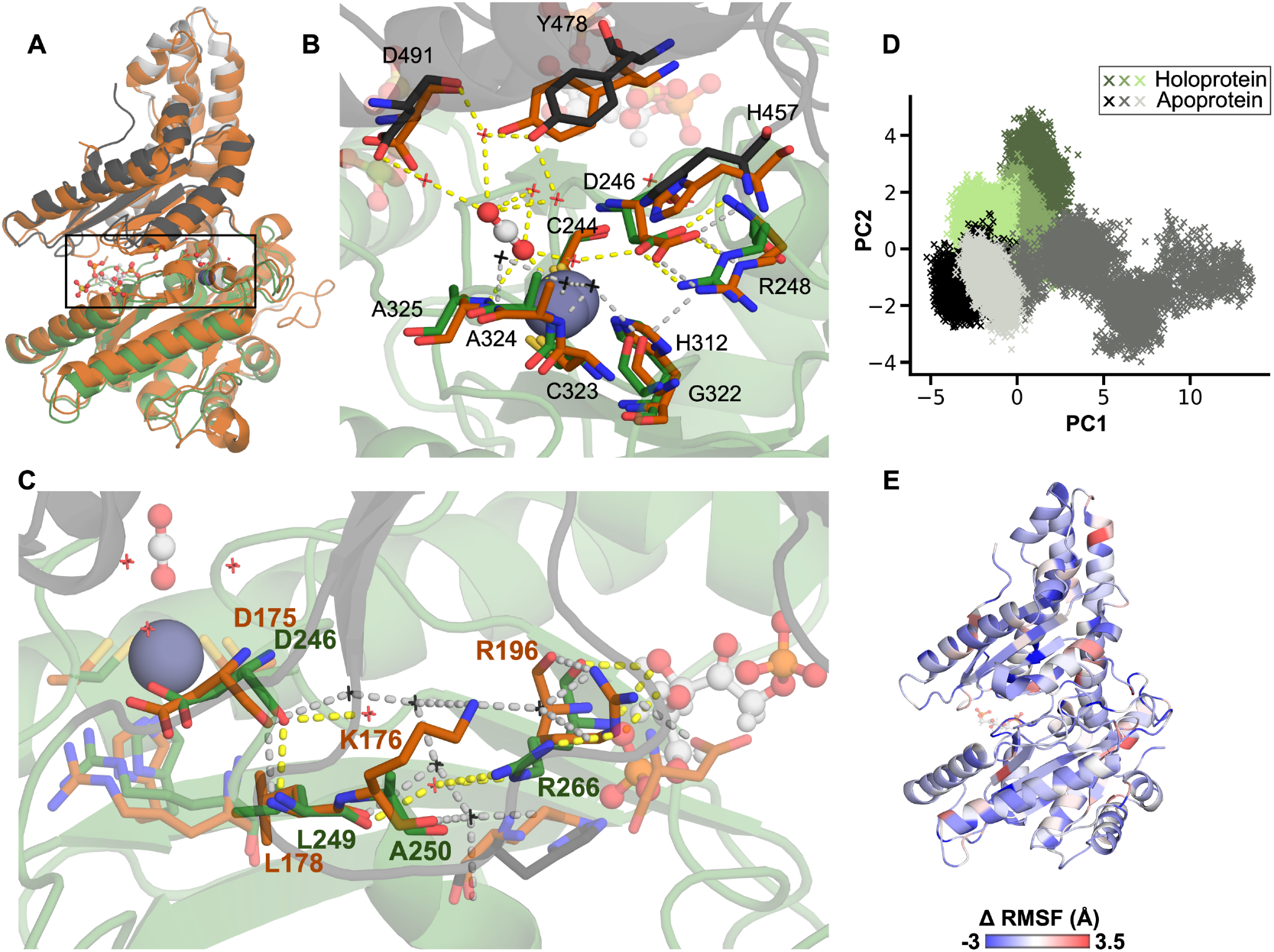
Active site differences from HnCsoSCA are key to allosteric activation of CyCsoSCA. **A** Structural alignment of CyCsoSCA and HnCsoSCA monomer (CA RMSD 1.5 Å). The CyCsoSCA variant is coloured by domain (white: N-terminal domain (NTD), green: Catalytic domain (Catalytic) and dark grey: C-terminal domain (CTD)). HnCsoSCA (PDB 2FGY) is shown in orange. A box denotes the active site and RuBP binding site region. **B** A close view of the CyCsoSCA active site is shown with key residues in stick representation, coloured as in A. Corresponding HnCsoSCA residues (orange) are overlaid. All ligands (CO_2_, RuBP, SO_4_) are shown in ball and stick representation. Catalytically relevant water molecules are shown as crosses (black waters are those in the HnCsoSCA structure, red indicates those in CyCsoSCA). Dashed lines indicate polar bonds (grey indicating those between HnCsoSCA molecules, yellow for CyCsoSCA interactions). Residue names are annotated according to the CyCsoSCA structure. **C** The proposed allosteric network in CyCsoSCA (top, green) and the corresponding region in HnCsoSCA aligned. Key residues for each structure are annotated in orange (HnCsoSCA) or green (CyCsoSCA). **D** Principal component analysis (PCA) comparing cartesian coordinates of the CyCsoSCA backbone in each MD simulation. Replicate simulations of CyCsoSCA with and without RuBP are shown in green (Holoprotein) and grey (Apoprotein), respectively. **E** The average difference in root mean square fluctuation (RMSF) values of holoprotein and apoprotein simulations (Δ RMSF), where a negative value indicates a greater RMSF (and thus more mobile residue) in the holoprotein. Values are mapped onto the CyCsoSCA monomer.

Molecular dynamic simulations of the *Cy*CsoSCA structure (300 ns replicates) were conducted in the presence (holoprotein) and absence (apoprotein) of RuBP to assess for conformational changes upon ligand binding to evaluate the allosteric effect of RuBP. Analyses of replicate simulations are consistent with *Cy*CsoSCA accessing different conformational landscapes when RuBP is present or absent. A principal component analysis of the trajectories highlights differences between conformations sampled in apo- and holoprotein states, with holoprotein replicates converging on a distinct cluster as the simulations equilibrate (Fig. 2D, S6). On average, residues in the apoprotein trajectories across the structure had root mean square fluctuations (RMSF), indicating they are more mobile (Fig. 2E). While it is difficult to ascribe a molecular mechanism with confidence, these results support a model in which RuBP stabilises *Cy*CsoSCA by establishing an internal H-bond network, promoting access to the active conformation.

### Sequence patterns indicate allosteric CsoSCA is limited to cyanobacteria

We sought to investigate the prevalence of RuBP allostery within the broader CsoSCA protein family by mapping the sequence diversity of the family to *Cy*CsoSCA functional variation. To examine CsoSCA divergence, a maximum likelihood phylogeny was inferred from a curated sequence database of the CsoSCA PFAM (PF08936) (Fig. 3A, S7). Cyanobacteria form a clear, tight cluster distinct from other bacterial species, supported by a high bootstrap value, as seen in related studies^33, 41, 42^. We hypothesised RuBP regulation may be specific to photoautotrophs, evolving as the *cso* operon adapted to these organisms’ light-dependent metabolic requirements relative to ancestral chemoautotrophic ɑ-carboxysomes^42^.

**Figure 3.**
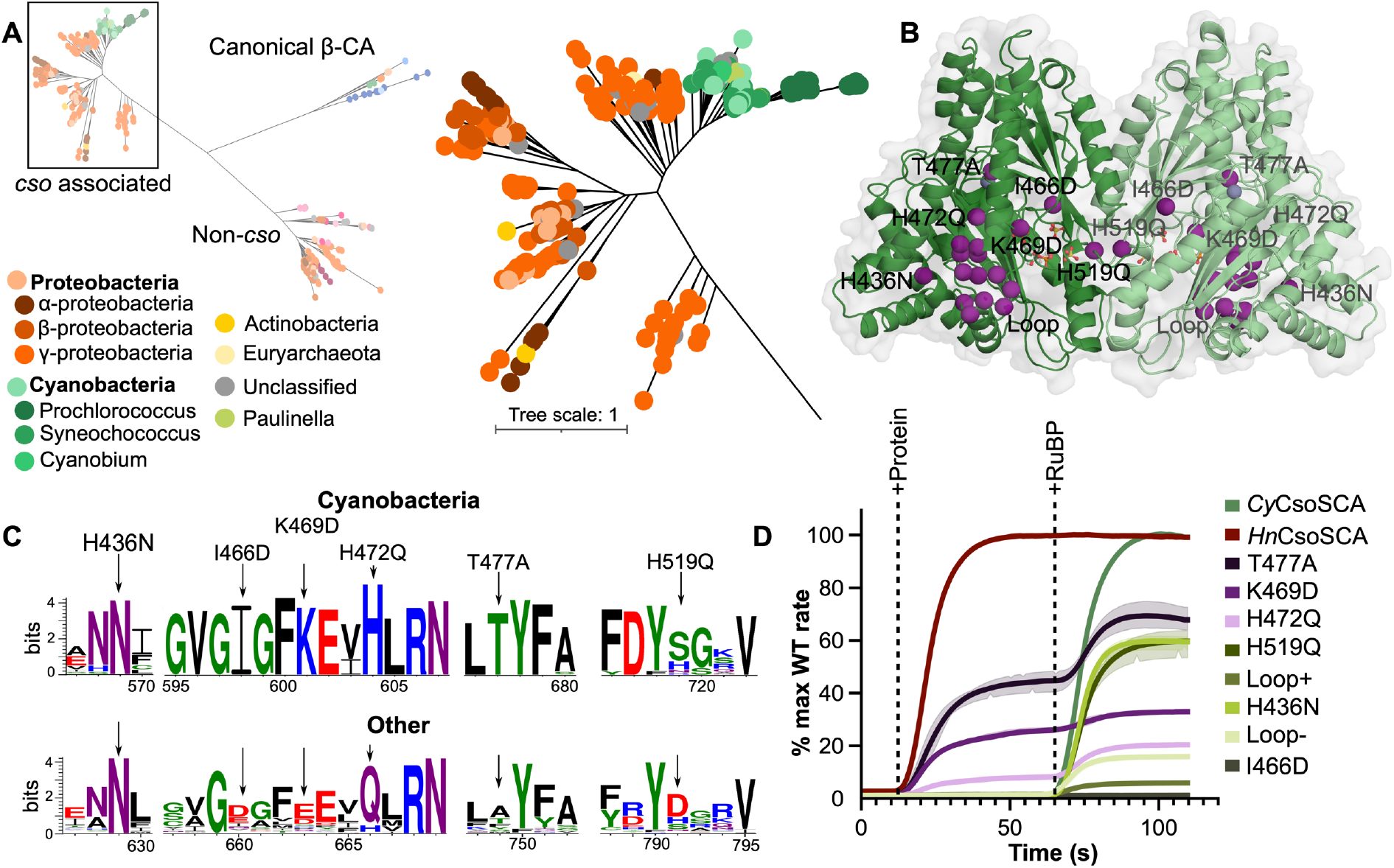
CsoSCA sequence analysis and mutagenesis. **A** Unrooted maximum likelihood phylogeny of 518 CsoSCA sequences produced by IQ-Tree with 18 canonical β-CAs as an outgroup (annotated as ‘β-CA’). The cluster containing members associated with the *cso* operon is shown in detail. Tips are coloured by taxonomy according to the legend. Tree scale refers to the number of substitutions per site. See supplementary information for complete tree annotation and documentation. **B** Sites targeted for mutagenesis shown as purple spheres on the *Cy*CsoSCA dimer. **C** Sequence logos based on cyanobacterial *cso* associated CsoSCA sequences (‘Cyano’) or other bacterial *cso* associated CsoSCA sequences (‘Other’) of key sites targeted for mutation. Residues coloured by chemistry, logo generated through WebLogo3. Mutations are as represented above the logo **D** Activity assays of targeted mutants relative to the maximum rate recorded for wild type *Cy*CsoSCA. *Hn*CsoSCA activity is also shown for comparison as a proportion of its maximum recorded rate. HCO_3_^-^ dehydration activity was recorded using MIMS after the addition of CsoSCA variants (+Protein) and upon addition of 100 µM RuBP (+RuBP). Three technical replicates were recorded for each variant, standard deviation is indicated by shading (error on some samples not visible due to scale).

To test this, concurrent approaches based on rational design and directed evolution were used to discern key residues involved in RuBP regulation. The conservation of these positions was then assessed across the CsoSCA protein family (Fig. 3). Candidate residues for targeted mutagenesis were chosen through successive steps of analysing the sequence and structure of *Cy*CsoSCA and *Hn*CsoSCA to locate residues with distinct biophysical properties near the RuBP pocket. Final mutations were made at sites that differed between the two characterised isoforms and had varying levels of conserved difference between cyanobacterial taxa and other bacterial CsoSCA isoforms more broadly (K469D, H472Q, I466D, H436N). Manual sequence inspection revealed a loop region in *Hn*CsoSCA (position in *Hn*CsoSCA) that contained an insertion conserved across cyanobacterial species (position in *Cy*CsoSCA) but absent or non-conserved in other proteins. Mutants were created in the *Cy*CsoSCA background with either a deletion of this loop (Loop deletion) or with the corresponding *Hn*CsoSCA loop sequence substituted at this site (Loop insertion). Alongside this approach, *Cy*CsoSCA was also randomly mutagenized, facilitating a broader exploration of the sequence space involved in this activation mechanism.

Mutants were screened using an in-house CA knock-out *E. coli* strain^43^ for variants with CA activity independent of RuBP. Activity assays of these mutants revealed that, in addition to K469D, a H472Q mutation (targeted approach) and T477A (random approach) also resulted in a biphasic activity profile with CA function independent of RuBP (Fig. 3D). Other mutations resulted in either reduced or undetectable CA activity, making their effects on RuBP dependence specifically, difficult to infer. Sequence-based analyses show residues with apparent involvement in RuBP-dependence are well conserved in cyanobacteria, but absent or non-conserved in other taxa (Fig. 3C). The conservation of sites underpinning RuBP-dependence is consistent with this regulation existing primarily, if not exclusively, in photoautotrophic CsoSCA variants.

### The unique N-terminal oligomerisation domain is exclusive to carboxysomal CAs

Further bioinformatic analysis of the CsoSCA protein family revealed an orphan cluster within this β-CA clade (Fig. 4). Manual inspection of the gene neighbourhoods of these sequences demonstrates they are not associated with the *cso* operon, appearing instead within other metabolic gene clusters, often associated with NADH or [Fe-Ni] hydrogenases or permeases (Fig. S12). Structural modelling of ‘non-*cso*’ sequences and subsequent structure-based searches using Foldseek^44^ and DALI^45^ shows these ‘non-*cso*’ sequences align preferentially to the published *Hn*CsoSCA structure with high confidence relative to other β-CAs (Table S4). These sequences appear to have lost the typical β-CA two-fold symmetry, containing only one predicted Zinc binding site per pseudo dimer, a defining feature of canonical CsoSCA sequences (Fig. 4). However, all ‘non-*cso*’ sequences are notably shorter than carboxysome associated variants. This is underlined by a consistent insertion in *cso-*associated sequences within the NTD which encodes a hook-like bundle of α-helices dubbed the ‘hook’ motif (Fig. 4) shown here to facilitate structural Zinc binding and oligomerization (Fig. 3). This is consistent with NTD presence, and thus hexamer formation being a more recent adaptation unique to carboxysome-encapsulated variants of this family.

**Figure 4.**
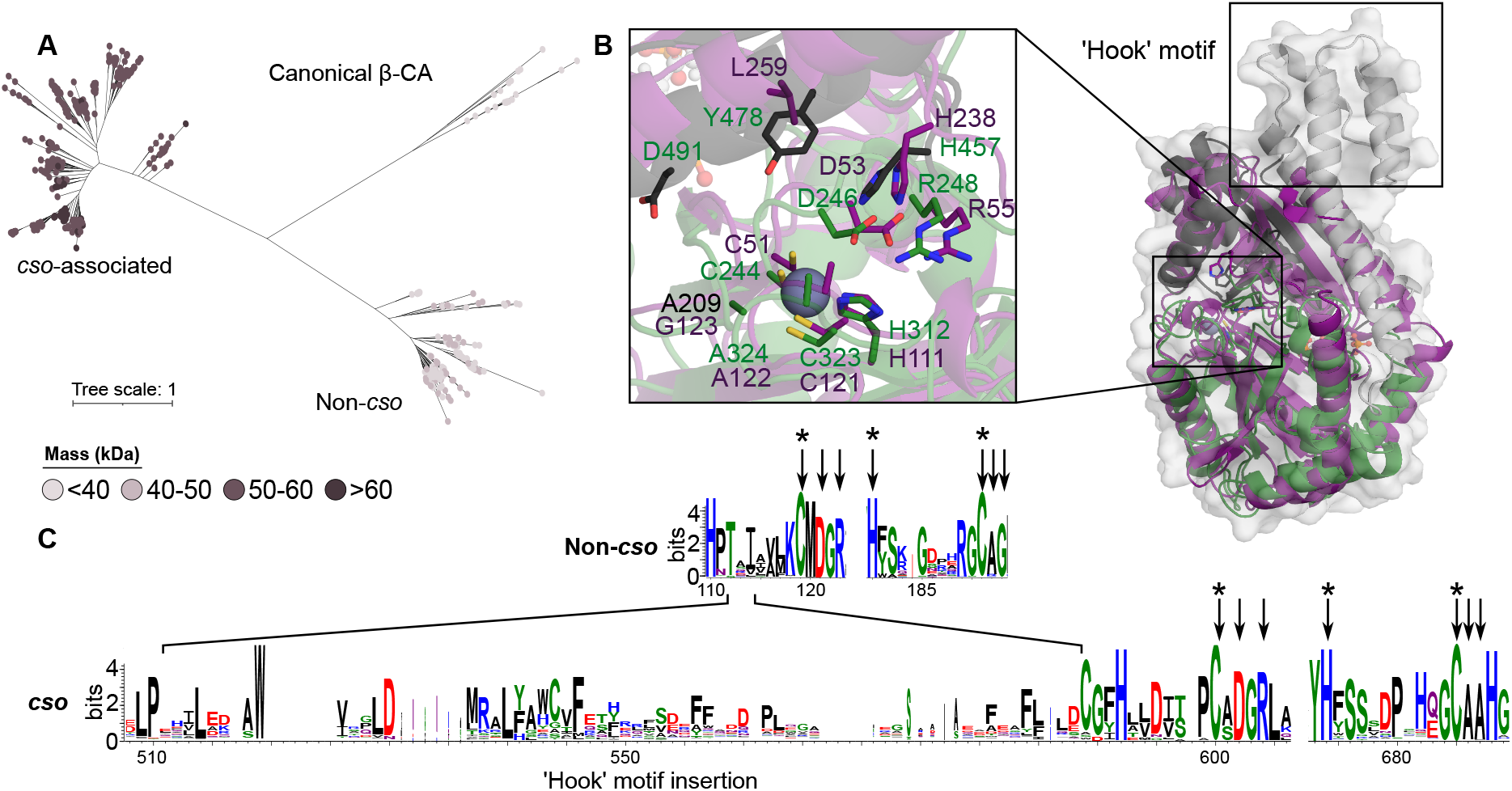
The N-terminal domain is unique to carboxysome operon-associated CsoSCA homologues. **A** Maximum likelihood tree of the CsoSCA PFAM (PF008936) and canonical β-CA with nodes coloured by predicted mass as per the legend. **B** Structural alignment of Cɑ backbones of *Cy*CsoSCA (NTD grey, Catalytic domain green, CTD black) to an AF2 generated model of a candidate ‘non-*cso’* sequence (UniProt ID: A0A080M7C6, purple) with close-up view of key active site residues. The insertion exclusive to *cso*-associated sequences that encodes the NTD ‘hook’ motif is annotated. **C** Sequence logo of alignments of either CsoSCA members associated with *cso* operons (*cso)* or non-*cso* sequences. The *cso* insertion encoding the NTD hook motif is annotated. Key catalytic residues are indicated with arrows, zinc binding residues with asterisks.

### Carboxysome functional modelling indicates an adaptive advantage for RuBP regulated CsoSCA

The results presented above support the hypothesis CsoSCA RuBP-dependence is a fixed trait unique to cyanobacterial α-carboxysome systems. To determine whether this feature emerged as an adaptive or neutral change in CsoSCA variants, we aimed to discern a functional benefit for RuBP regulation within the cyanobacterial system. An *in vivo* assessment of CsoSCA regulation is currently intractable, with no effective genetic transformation techniques reported to date. Instead, we modified our carboxysome steady state diffusion model^38^, incorporating kinetic data presented in Figure 1A to compare the activity of a *Cyanobium* carboxysome with and without an RuBP-dependent CA (Fig. 5A). While second to physiological data, this approach permitted insights into the effects this regulation may have on core enzyme activity and metabolite flux of the *Cyanobium* α-carboxysome. No substantial changes in Rubisco carboxylation or oxygenation rates were observed between the standard model and one incorporating an RuBP-dependent CA (Fig. 5B, S9). However, we noted that the modelling of the unmodified *Cyanobium* carboxysome here had a slightly alkaline carboxysome pH compared with the observation of an acidic carboxysome in our previous modelling^38^. This is due to the use of *Cyanobium* Rubisco kinetics, the concentration of HCO_3_^-^ tested here (20 mM), and a modification of the Rubisco active site concentrations compared with previous modelling (a reduction from 10 mM to 5.7 mM to fit recent estimates^32^). However, applying RuBP-dependence on CA activity in the model provides a shift to a more acidic condition that can support Rubisco carboxylation^38^ and potentially ensures Rubisco is maintained within its optimum pH range for activity^46^. This implicates a regulated CA in maintaining an acidic carboxysome lumen under low RuBP conditions in the *Cyanobium* system, likely experienced as light levels fluctuate throughout the diurnal cycle^20^, thereby contributing to efficient CCM function^47^.

**Figure 5.**
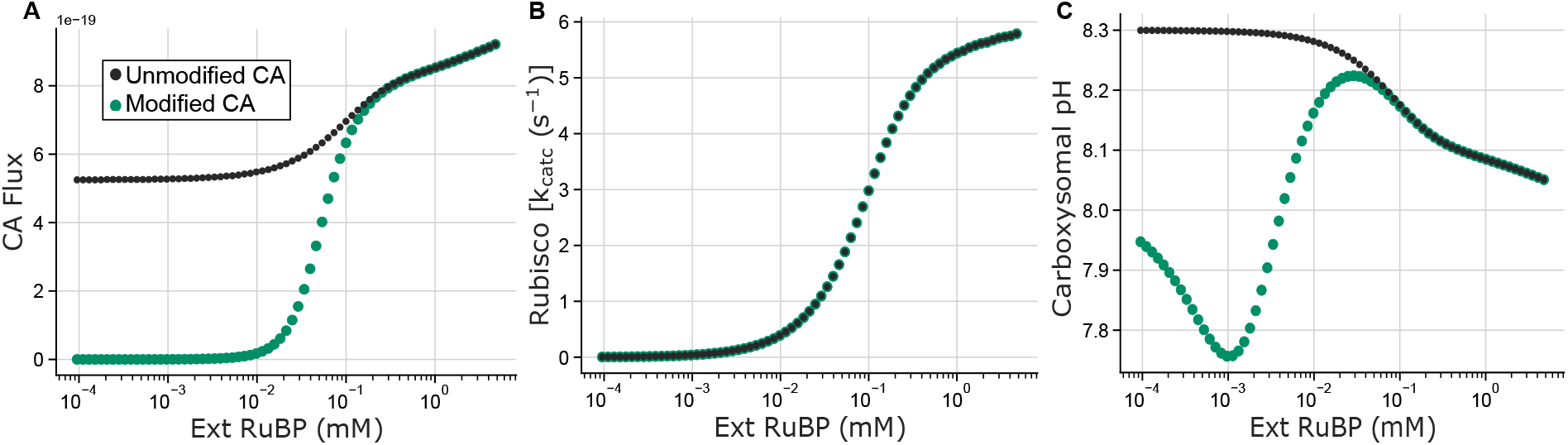
Results of a reaction-diffusion model^38^ adapted to emulate *Cyanobium* α-carboxysome function with and RuBP d-dependent (Modified CA, green dots) or a constituently active (Unmodified CA, black dots) CA. **A** In the modified model, carboxysomal CA activity was altered based on data shown in Figure 1A to be dependent on carboxyosmal RuBP concentrations (Ext RuBP (mM)). ‘CA flux’ is indicative of CA activity, confirming RuBP dependence in the modified model. **B** Rubisco carboxylation turnover rates (Rubisco [*k_catc_* (s^-1^)]) as a function of modelled cellular RuBP concentrations (Ext RuBP (mM)) in the modified and unmodified systems. **C** Modelled carboxysomal pH is plotted as a function of the modelled cellular RuBP content (Ext RuBP (mM)) indicates a decrease in carboxysomal pH when CA function is allosterically controlled by RuBP, with potential to aid Rubisco function as described previously^38^.

## DISCUSSION

The ɑ-carboxysomal CCM enables efficient C_i_ fixation across a diverse range of microorganisms, comprising a major component of the global biosphere^5, 48^. Given the essential nature of CAs in this system^6^, characterising the CsoSCA variant present in photosynthetic organisms, and understanding its evolutionary trajectory provides important insights into the emergence and function of bacterial CCMs. Data presented here are consistent with RuBP allosterically activating *Cy*CsoSCA and the likely confinement of this property to α-cyanobacteria. A hexameric ‘trimer of dimers’ quaternary state is also described, coordinated by NTD contacts with structural zinc ions that appear only in carboxysome-associated members of the CsoSCA protein family.

### A distinct paradigm for carbonic anhydrase allosteric regulation

To our knowledge, this is the first case of allosteric activation reported for a carbonic anhydrase. The β-CA family is the only CA family known to exhibit allostery, with HCO_3_^-^ acting as both a substrate and inhibitor in Type II members. In that case, HCO_3_^-^ binding disrupts the ‘gate-keeper’ Asp-Arg dyad, leading to Asp-Zn bond that displaces the catalytic H_2_O, leading to inhibition^40, 49–51^. In addition to *activating* the enzyme, the RuBP binding pocket presented here appears distinct from these previously characterised sites, engaging different residues and sitting further from relative active sites (Fig. S4). Indeed, the RuBP site sits near the defunct active site of the CTD within the CsoSCA pseudo-dimer, previously identified as a highly divergent catalytic domain that has lost key zinc-binding sites and catalytic loops^30^. The duplication and divergence of domains, particularly at protein termini, are a common motif in protein evolution^52^. While seemingly acting as a regulatory domain in the cyanobacterial CsoSCA, it is unclear whether the CTD takes on alternative, more cryptic regulatory roles in RuBP-independent CsoSCA isoforms, or indeed in the smaller uncharacterised ‘non-*cso*’ isoforms.

We propose an allosteric activation mechanism in which RuBP binding establishes an internal H-bond network near the core of *Cy*CsoSCA that has a broadly stabilising effect on the protein, this in turn promotes access to the active conformation. RuBP binding engages Arg266 in H-bonding, establishing a H-bonding network that links to active site loops (Fig. 3). The equivalent network in HnCsoSCA is more extensive involving more residues, specifically negatively charged residues that are absent in the CyCsoSCA isoform. Specifically, in *Hn*CsoSCA, the Arg266 equivalent (Arg196) is stabilised by analogous interactions with Asp409 (Lys469 in *Cy*CsoSCA), thereby negating the ligand-binding requirement for activation. This proposed mechanism is also consistent with the observed biphasic activity profile of mutants many of which introduce a relatively negative charge to this site, most notably K469D (Fig. 3). However, it remains difficult to rationalise the relatively large increase in RuBP-independent activity observed in the *Cy*CsoSCA^T477A^ mutant. While Thr residues are typically known to stabilise β-sheets, given this site is buried it may be that this exchange reduces secondary structure strain and enhances protein core packing while with more minimal disruption to the overarching protein fold that other mutants^53^. Consequently, T477A would result in increased apoprotein stability *and* a WT-like RuBP pocket, leading to high levels of ligand-independent activity and a notable increase with RuBP. Although the involvement of the Asp-Arg dyad destabilised by HCO_3_^-^ does seem reasonable in the context of *Cy*CsoSCA given it’s established role as an allosteric switch^40, 49^, we could not establish this as the clear mechanism for allosteric signal propagation.

### An adaptive advantage for an RuBP-regulated CA in photoautotrophs

The conservation of residues central to RuBP-dependent activity corresponds with a clear divergence of the cyanobacterial clade in the CsoSCA phylogeny (Fig. 3). This suggests allosteric RuBP activation is either a neutral or adaptive change uniquely fixed in ɑ-cyanobacteria relative to other ɑ-carboxysomal taxa. RuBP allostery appears infrequently in the literature, primarily characterised in Rubisco adjacent proteins such as the AAA+ red-type activase Cbbx^54^. In these cases, it is proposed to synchronise the activity of such proteins with Rubisco, hinting at an intricate regulatory cycle to ensure efficient Rubisco function. Indeed, modelling and experimental data have established a similar functional link between CA and Rubisco, demonstrating Rubisco function in carboxysomes is intrinsically dependent upon CA activity^6, 28, 38^. This type of post-translational regulation would enable more rapid responses to transient metabolic signals, directly synchronising Calvin cycle fluxes with Rubisco-mediated carbon fixation.

We propose RuBP-dependent allostery may be linked to large fluctuation in cellular RuBP observed in photosynthetic ɑ-cyanobacteria, but absent in proteobacterial systems such as *H. neapolitanus* that are likely under more constant environmental substrate supply^20, 55^. Using a carboxysome reaction-diffusion model, we previously identified that carboxysomes may require molecular mechanisms that modulate internal pH as RuBP concentrations vary, to the detriment of Rubisco function^38^. Modification of this model to allow for allosteric activation of CA in *Cyanobium*-like carboxysomes indicates such regulation ameliorates this effect, resulting in a modulation of carboxysome pH without any change in Rubisco carboxylation (Fig. 5, S9). Thus, the emergence of RuBP regulation in these systems may have been prompted by a requirement for more fine-tuned control over carboxysomal H^+^ concentrations in photoautotrophic systems that are subject to more drastic cellular RuBP fluctuations across the diurnal cycle. That the protein can be switched from a constitutive to autoinhibitory isoform in one residue change highlights the evolutionary pliability of this protein, suggesting the evolution of allostery from a constitutively active isoform may have arisen from a relatively small sequence level change in ancestral photoautotrophic hosts in the presence of new fitness pressures. This raises the question as to why β-carboxysomes, present exclusively in photosynthetic cyanobacteria, does not appear to have the same requirement?^56, 57^ Our modelling has previously identified that the larger size of β-carboxysomes, and thereby surface area:volume ratio, relative to ɑ-carboxysomes leads to diffusional differences between these structures^38^. In this way, β-carboxysomes may not be as vulnerable to the internal species fluctuations that underpin the proposed advantage of an RuBP-regulated CA. Given this, we conclude the emergence of an RuBP-dependent CA comprises a key molecular step in the adaptation of the ɑ-carboxysome within cyanobacterial lineages.

### The ɑ-carboxysomal CA is hexameric in solution

The oligomeric state of enzyme cargo is an important detail of bacterial microcompartment systems, providing insights into cargo organisation and interaction networks within the shell. Here, we show that both *Cy*CsoSCA and *Hn*CsoSCA are both hexameric in solution, not dimeric as previously described^30^, and that this quaternary structure is mediated by the previously unannotated NTD (Fig. 1). This hexamer likely eluded detection in the reported *Hn*CsoSCA structure due to an inadvertent mutation causing an artificial second zinc site within the NTD of each monomer that would likely inhibit the structural zinc interactions observed here^30^ (Fig. S11). Given this mutant still exhibited high levels of CA activity, it is unlikely the two additional zinc sites detailed here are catalytically relevant, instead acting primarily to stabilise the quaternary structure. Moreover, the analysis of the CsoSCA protein family presented here suggests the NTD, and thus the ability to form this quaternary structure, is specific to carboxysome-associated CsoSCAs (Fig. 4). CsoSCA is now known to interact with Form IA Rubisco through interactions between the CsoSCA N-terminal disordered tail in a manner reminiscent of the ɑ-carboxysome structural protein CsoS2^33, 57^. Though the conservation of this binding motif is limited, often each N-terminal disordered tail appears to be limited to a single motif. Comparatively, CsoS2 contains multiple Rubisco binding motifs, enhancing the multivalency and thus strength of the interaction. Considering this, a hexameric assembly would enhance the local concentration of Rubisco interaction motifs and thus multivalency of the system to promote CsoSCA-Rubisco binding. In this way, the emergence of the NTD may constitute a molecular marker for the early association of an ancestral CsoSCA-Rubisco complex, hypothesised as a likely carboxysome evolutionary precursor^38, 58^.

## CONCLUSION

As we begin to resolve the evolutionary trajectories of bacterial CCMs, a detailed understanding of structural variation of core system components and how this relates to function will be essential for resolving plausible evolutionary routes. We have identified a distinct divergence between α-cyanobacteria and other α-carboxysome taxa, presenting a paradigm for CO_2_ fixation in photoautotrophic α-cyanobacteria that hinges on the regulation of CsoSCA by RuBP. These results highlight the intrinsic role of CAs in photosynthesis, comprising an evolutionarily pliable enzyme at the bottleneck of key reactions in the carbon metabolism of many diverse organisms. This aspect of carboxysome function and evolution must be considered in future biotechnological applications that seek to adapt such systems, particularly efforts to transplant the bacterial CCM into photoautotrophic crop species.

## Acknowledgements

The authors acknowledge use of the Australian Synchrotron MX facility and thank the staff for their support. This research was undertaken in part using the MX2 beamline at the Australian Synchrotron, part of ANSTO, and made use of the Australian Cancer Research Foundation (ACRF) detector.

## Funding

Funding for some salary components (BF, SBP, BML), materials, DNA constructs, and sequencing work was supported by a grant from the Australian Research Council Centre of Excellence for translational Photosynthesis (CE140100015) to GDP. CJJ and SBP also acknowledge support from the Australian Research Council Centre of Excellence in Synthetic Biology (project number CE200100029). SBP acknowledges stipend funding from the Westpac Scholars Trust.

## Author contributions

SBP and BML conceptualisation, experimental design, data collection and analysis, writing, editing; MAO Data collection and analysis; BF Data collection; SJW Data collection and analysis; CJJ Data collection and analysis, writing and editing, funding; MRB conceptualisation, experimental design; GDP conceptualisation, experimental design, editing, funding; TR experimental design, data collection.

## Competing interests

Authors declare that they have no competing interests.

## Data and materials availability

All data are available in the main text or the supplementary materials. The *Cyanobium* CsoSCA structure is available at the PDB ID: 8THM.

## Notes

### Competing Interest Statement

The authors have declared no competing interest.

