## Supplemental Materials for "Cyanobacterial α-carboxysome carbonic anhydrase is allosterically regulated by the Rubisco substrate RuBP"

**This PDF file includes:**

Materials and Methods
Supplemental Methods
Tables S1to S5
Figs. S1 to S12
Supplementary References

**Other Supplementary Materials for this manuscript include the following:**

Data S1: Master csv of sequences used in analyses
Data S2: Treefile for final inferred CsoSCA phylogeny

### Materials and Methods

CsoSCA expression constructs

In initial expression tests, we found the N-terminal disordered region (Fig S1) causes severe aggregation, inhibiting expression and purification and thus was removed in all CsoSCA isoforms discussed in this investigation. The globular regions of wild type *Hn*CsoSCA and *Cy*CsoSCA were codon-optimised for expression in *E.coli* and cloned into the pHue expression system[59]. Codon-optimised genes encoding *Cy*CsoSCA mutants were synthesised through Twist Bioscience and cloned into the pHue vector using standard Gibson assembly methods[60]. Primers used are listed in Table S5.

CsoSCA random mutagenesis

To create random CsoSCA mutants with RuBP-independent activity profiles, random base changes were introduced using error prone pCR (epPCR) and the resulting libraries were screened for desired activity profiles. The GeneMorph II kit (Agilent, Cat. No. 200550), containing the Mutazyme II DNA polymerase blend, was used in the epPCR reaction to generate mutant libraries, using conditions specified in the manual for medium rates of mutagenesis. Primers were designed to enable a one-pot cloning reaction (Engler et al., 2008) of resulting PCR fragments into a pET16 expression vector (Table S5). The resulting one-pot reaction mix was dialysed using MF-Millipore 0.25 µm MCE Membrane (Merk, Ireland) to remove salts and transformed into dense aliquots of electrocompetent DH5α *E. coli* cells. The library was plated and grown overnight at 22°C until pinprick colonies were visible. Plates were then washed with 5mL LB to recover colonies and plasmids extracted using the QIAprep Spin Miniprep Kit (Qiagen, Cat. No. 27104). The DCAKO cell line is an *E. coli* strain, created in Price lab, in which both native CA genes have been knocked out. Consequently, cells cannot grow under ambient conditions, likely due to insufficient cytosolic $HCO_3^-$ concentrations required for the central carbon metabolism (Merlin et al., 2003). 5µL of this library was transformed into dense aliquots of electrocompetent DCAKO E. coli cells to screen the mutant library for members with the desired activity profile (activity independent of RuBP). Transformed cells were grown in triplicate plates at 37°C/ambient or 37°C/4% $CO_2$ conditions. Colonies that appeared on plates grown under ambient conditions were re-grown in liquid cultures at 37 °C/4% $CO_2$ and plasmids extracted for sequencing. Resulting mutant *csoSCA* sequences analysed using Geneious Prime software package. Mutants that permitted DCAKO growth under atmospheric conditions were sequenced and a representative pool was chosen for crude activity assays to confirm CsoSCA activity independent of RuBP. Three residues (H519Q, T477A, N278D) consistently corresponded to RuBP independent CA activity, all had notably different chemistry to the analogous sites in *Hn*CsoSCA. However, in the mutant library, these sites were always present within a small background of other site changes. To assess the effect of these residues directly, single site mutants were generated using custom primers and cloned into the pHue expression system (Table S5). The N278D mutant was incredibly difficult to clone and consistently precipitated during expression leading us to abandon it for further analysis.

<u>Protein expression and purification</u>

The USP2-pHue expression system described by Catanzariti et al. (2004) was employed for heterologous expression of CsoSCAs[59]. NEB T7 Express pLysY chemically competent *Escherichia coli* cells were transformed with a pHue plasmid containing the corresponding CsoSCA sequence. Single colony glycerol stocks (1:1 ratio of 40% glycerol and cell culture) were used to inoculate 3ml cultures in Luria-Bertani (LB) media supplemented with 100 μg/mL ampicillin, and grown overnight at 37°C shaking at 200 RPM. From these overnight cultures, 1ml was used to inoculate 25ml fresh LB+Amp which in turn was incubated at 37°C for 6 hours in LB media, 10ml of this was then used to inoculate 500ml fresh LB+Amp. This was grown at 37 °C for 2 hours, followed by induction with 100 μM isopropyl-β-D-thiogalactopyranoside (IPTG) and further incubation overnight at 28°C. Cell cultures were pelleted and stored at -80°C until required.

Cells were thawed, resuspended in binding buffer (50 mM Tris [pH 7.8], 300 mM NaCl, 25 mM Imidazole) and incubated at room temperature shaking with 10% rLysozyme (EMD Millipore Corp, USA) and 0.2 μl Turbonuclease (Sigma Aldrich) for 30 minutes. Cells were lysed with three passes of the Emulsiflex (Avestin, USA). Lysate was clarified by centrifugation (16,000xg for 30 min at 4°C) and the soluble fraction passed through a 0.44 μm syringe filter before application to a pre-equilibrated 5 mL HisTrap HP column (GE Healthcare). The column was washed with 50 mL binding buffer and the target protein eluted in elution buffer (50 mM Tris [pH 7.8], 300 mM NaCl, 500 mM Imidazole). Fractions containing protein were pooled and concentrated to a maximum of 5 mL using a centrifuge filter (Amico Ultra-15 Centrifugal Filter Unit) and buffer exchanged using a PD-10 column (GE Healthcare, Lot 9760001) into Size-Exclusion Buffer (SEC buffer, 50 mM Tris [pH 7.8], 300 mM NaCl). Samples were incubated with pre-purified USP2 at a 1:10 protein-of-interest:USP2 molar ratio and incubated rocking at 4C for at least 12 hours with 2μL βmercaptoethanol. Samples were then passed over clean HisTrap columns equilibrated with 50ml SEC buffer to remove protease. Flow through was collated and concentrated through centrifugation as above and further purified by size exclusion chromatography on a HiLoad 26/600 Superdex 200 Column (GE Healthcare), eluting in SEC Buffer. Protein purity was confirmed by SDS-PAGE, and protein concentrations were determined spectrophotometrically by measuring A280 using a NanoDrop One (Thermo Scientific) and molar absorption coefficients generated by ProtParam (http://expasy.org/tools/protparam.html).[61]

#### Carbonic anhydrase activity assay

All carbonic anhydrase (CA) activity was measured using the Membrane Inlet Mass Spectrometry (MIMS) technique and relevant equations detailed in Badger and Price (1989) adapted for pure protein samples[29]. This is based on measuring the loss of $^{18}O_2$ from a labelled $C_i$ source to water through the CA catalysed hydration and dehydration of $CO_2$ and $HCO_3$-, respectively. Briefly, $NaH^{13}CO_3$ is incubated for 24 hours with $H_2{}^{18}O$, creating a $^{13}$C-labelled carbon source enriched with $^{18}$O. The change in the $^{18}$O enrichment of $CO_2$ species is measured after chemical equilibrium was reached before and after the addition of enzyme. The $^{18}$O atom fraction (the atom % enrichment) was calculated at each point by summing the related Carbon species determined by Mass Spectrometry. CA activity is reported as the rate of decline of the atom % enrichment, calculated from the slope of the change in log(atom % enrichment) over time ($Log_{Enrich}$ $min^{-1}$). This gives a measure of the first order rate constant. As this refers to the decline of $^{18}$O, this rate becomes increasingly negative with CA activity. For simplicity we display the absolute value here. All assays were conducted with 0.1uM protein and 2.7 mM heavy isotope bicarbonate ($NaH^{13}C^{18}O_3$) in CA buffer (50mM EPPS-NaOH, 20mM MgCl2 [pH 7.8]) in a final cuvette volume of 600 µL. To standardise oxidation of samples, 30 µM 5,5'-dithiobis-(2-nitrobenzoic acid) (DTNB) was added to reaction mixes before measurement. Final cuvette volume was brought to 600µL. The rate of consumption of each measured isotope was determined at equilibrium before and after protein was added to the system and upon addition of RuBP when relevant. Activity measurements for *Cy*CsoSCA mutants were conducted as above all in the presence of 100µM RuBP.

#### Crystallisation, data collection and structure determination

Initial screening to determine crystallisation conditions was performed at *Cy*CsoSCA concentrations of 10mg/ml at 20/18 °C in 96 well plates using the sitting drop vapour diffusion method and commercially available sparse matrix screens (ShotGun SG1 (Molecular Dimensions), PegIon (Hampton), Crystal Screen HT (Hampton), JCSG (Molecular

Dimensions), PACT Premier (Molecular Dimensions), Index HT (Hampton), PegRx (Hampton), Salt Rx (Hampton)). In each case, 200nl drops comprising 100nl protein solution and 100nl reservoir were prepared on hanging-drop seals using a NT8®-Drop Setter robot (Formulatrix, Bedford, MA, USA) and equilibrated against 100 µl reservoir solution. Further optimisation was carried out in 24-well hanging-drop vapour diffusion plate format varying pH, precipitant and protein concentration conditions. The final optimised crystallisation conditions were at 0.2M Ammonium sulfate, 0.1M BIS-TRIS (pH 6.5), 21% PEG 3350 and 15% Ethylene glycol, seeded with crystals from the same conditions at a protein concentration of 7mg/mL and an RuBP:CyCsoSCA molar concentration of 10:1. Crystals were cryoprotected with glycerol and flash-cooled in liquid nitrogen. X-ray diffraction data were collected at beamline MX2 at The Australian Synchrotron. Data was processed using XDS and Aimless, and molecular replacement was performed using Phaser. The ColabFold: AlphaFold2 using MMseqs Google Colab notebook to generate a model of *Cy*CsoSCA for molecular replacement[62,63]. Iterative cycles of manual model building and refinement were performed using Coot[64], ccp4[65] and *phenix.refine*[66]. TLS parameter refinement was also used with TLS groups automatically selected by *phenix.refine*. Data collection and refinement statistics are provided in Table S1. Structures were subsequently annotated and analysed in PyMOL.

Molecular Dynamics

The structure of *Cy*CsoSCA chain D reported here was used for all simulations. For holoprotein preparation Chain D was used as is, otherwise apoprotein starting structures were generated by deleting RuBP from the starting file. Automated structure refinement was conducted by the protein preparation wizard module of Maestro (Schrodinger, version 12.9.137 release 2021-3). This comprised assignment of bond order, addition of missing hydrogens, and generation of het states using Epik program at a target pH of 7.4. The hydrogen bond network was subsequently optimised with the default parameters and protonation states were assigned to residues at a pH of 7.4 using PROPKA. System builder was used to set up an orthorhombic box filled with TIP5P water models and the charge neutralised with 18 $Na^+$ ions, a buffer distance of 15Å was employed and the OPLS4 force field applied. A short minimisation was performed on the resultant structure of 1ns. Simulations were performed at a total time of 300ns at a time step of 50ps in three replicates for each system at different initial random seeds. Resultant trajectories were analysed using the Simulation Interaction Diagram (SID) module in Schröedinger and using the MDTraj python package[67].

Phylogenetic reconstruction

All CsoSCA sequences within the protein family (PF08936) were harvested from the pfam database. Duplicates were removed and a local allvsall BLAST search[68] was conducted to generate a sequence similarity network of the sequences with e-value cut-off of $10e^{-10}$. This was used to identify classes within the family and, using incremental percent identity edge cutoffs in Cytoscape (version 3.9.1), clusters within these classes were examined. An additional 20 reviewed sequences of known β-CAs were appended to this database for comparison with the 'non-*cso*' clade. Sequences without key catalytic sites and outliers were manually removed and the remaining sequences were filtered using CD-HIT to remove redundancy to 90%, resulting in a final database of 504 sequences, details of these, including accessions, are documents in SI (20 reviewed β-CAs, 484 CsoSCA protein family sequences)[69]. A multiple sequence alignment of these was constructed using MAFFT-LINSI[70]. Columns with >90% gaps were removed using TrimAI through the pheylemon2 web server (http://phylemon2.bioinfo.cipf.es/)[71]. This edited alignment was submitted to IQ-TREE (http://iqtree.cibiv.univie.ac.at/) for phylogenetic inference via maximum likelihood (ML)[72,73]. The ML sequence evolution model (WAG+F+I+G4) was selected as implemented in ModelFinder. Branch supports were measured by ultrafast bootstrap approximation and approximate likelihood ratio test, each conducted to 1000 replicates in IQ-TREE. Five independent replicates of tree-search were conducted, one of which was selected due to high ranch supports at key bifurcations. Trees were visualised and edited using iTOL (https://itol.embl.de/)[74]. Sequence Logos of alignments presented in the text were generated through WebLogo3 (https://weblogo.threeplusone.com/create.cgi)[75]. To assess the genetic context of non-*cso* sequences, the genetic context was visualised and manually inspected through JGI IMG (https://img.jgi.doe.gov/)[76].

COPASI Modelling

Modelling of carboxysome function with and without RuBP-dependent CA activity was carried out using the COPASI Biochemical Modelling Simulator as described by Long et al. (2021)[38]. Modelling was carried out for a single *Cyanobium* carboxysome of 100 nm diameter, containing a Rubisco active site concentration of 5.68 mM based on average number of Rubisco holoenzymes per Cyanobium carboxysome reported at 224 by Ni et al (2022)[32] and the Rubisco catalytic parameters reported in Long et al. (2021) ($kcat_C$ = 9.4 s-1, $K_{MCO2}$ = 169 µM, $K_{MO2}$ = 1.4 mM, $kcat_O$ = 1.42 s-1, $K_{MRuBP}$ = 40 µM)[32,38]. Modelling was carried out at 20 mM $HCO_3^-$, pH 8.0, and atmospheric $O_2$ concentrations over an exponential range of RuBP concentrations from 0.1 µM – 5 mM, spanning the apparent $K_M$ for CA activation (18 µM). Permeabilities were set to simulate a carboxysome (Long et al. 2021), with CA activity confined to the Rubisco compartment. The model was run with either unmodified CA function or modified to be dependent upon RuBP concentrations to replicate the observed RuBP response in Figure 1. Specifically, the model modifies CA activity as a function of the deprotonated, and most abundant, form of RuBP within the carboxysome ($RuBP^{4-}$).

Rate constant for the unmodified CA forward reaction ($CO_2 + H_2O \rightarrow HCO_3^- + H^+$):

$$CAfactor \times k_1 \times [CO_2]_c$$

Rate constant for the unmodified CA backward reaction ($HCO_3^- + H^+ \rightarrow CO_2 + H_2O$):

$$CAfactor \times k_2 \times [HCO_3^-]_c \times [H^+]_c$$

where the carboxysomal *CAfactor* is set to 100,000, $k_1$ is 0.05, $k_2$ is 100, and $[CO_2]_c$, $[HCO_3^-]_c$, and $[H^+]_c$ are the carboxysomal concentrations of $CO_2$, $HCO_3^-$ and protons, respectively[38]. Where CA function is not present in the model (in the external and unstirred compartments), *CAfactor* is set to 1 (1 × the background rate of interconversion).

Carboxysomal CA function within the model was modified to be dependent on RuBP concentrations by multiplying each of the forward and backward rate constants by the following function:

$$\frac{{[RuBP]_c}^h}{Khalf^h + {[RuBP]_c}^h}$$

where; $[RuBP]_c$ is the carboxysomal concentration of $RuBP^{4-}$, $h$ is the Hill slope (set to 2.214), and *Khalf* is the apparent $K_{MRuBP}$ required to achieve half maximal CA activity (18 µM). Both $h$ and $K_{half}$ were determined through Hill-reaction curve fitting of the RuBP response curve of *Cyanobium* CA in Figure 1, using Graphpad Prism.

### Supplemental Methods

Ion inhibition assays

To determine the effect of sulfate ions on CsoSCA activity, the standard buffer described above was substituted for either EPPS only (50mM EPPS-NaOH pH 7.8), 50mM MgCl2 (50mM EPPS-NaOH pH 7.8, 50mM MgCl2), 20mM MgSO4 (50mM EPPS-NaOH pH 7.8, 20mM MgSO4) and 50mM MgSO4 (50mM EPPS-NaOH pH 7.8, 50mM MgSO4). Measurements for *Cy*CsoSCA were taken in the presence of 100uM RuBP, given no dependency on RuBP was observed for *Hn*CsoSCA it was not added to these activity conditions.

Analytical SEC

A HiLoad 16/600 Superose 6pg preparative SEC column was calibrated using the Cytiva Gel Filtration Calibration HMW kit (product code 28403842) as per manual specifications (28951560 AG). The partition coefficient (Kav) of each protein was used to construct a calibration curve of Kav versus log(molecular mass). This was calculated using the equation Kav = (ve – vo)/(vc – vo), where ve is the elution volume, vo is the column void volume, and vc is the geometric column volume. These were used for comparison of elution columns and estimation of theoretical molecular masses.

Native PAGE

Protein samples were stored in Native Gel-Loading buffer (2.5 x TBE, 50% Glycerol, 0.1% Bromophenol blue) and loaded on 4–20% Mini-PROTEAN TGX Stain-free polyacrylamide gels (Bio-Rad, Cat. No. 4568096). Proteins were separated at 90 V for 90 minutes at $4^{o}$C in native running buffer (25 mM Tris [pH 8.3], 50 mM glycine). NativeMark protein standards were used for molecular weight determination (Invitrogen, LOT 1739010). To visualise proteins, gels were stained with Coomassie Blue dye (BioRad, USA). To determine how essential the observed zinc ions in the *Cy*CsoSCA structure are for hexamer formation, the protein was treated with a range of conditions to interrupt zinc binding, the oligomeric state of resulting samples was then assessed by Native PAGE. CyCsoSCA was dialysed with 2mM 1,10-phenanthroline, a strong chelating agent, for 24 hours according to previous publications[22]. Additionally, protein samples were incubated in SEC buffer at a pH of 3.5, 4, 5, 6, 7, or 8 for 1 hour. All samples were stored in Native Gel-Loading buffer (2.5 x TBE, 50% Glycerol, 0.1% Bromophenol blue) and loaded on 4–20% Mini-PROTEAN TGX Stain-free polyacrylamide gels (Bio-Rad, Cat. No. 4568096). Proteins were separated at 90 V for 90 minutes at 4°C in native running buffer (25 mM Tris [pH 8.3], 50 mM glycine). NativeMark protein standards were used for molecular weight determination (Invitrogen, LOT 1739010). To visualise proteins, gels were stained with Coomassie Blue dye (BioRad, USA).

```
52_Cyanobium_sp._PCC_7001  ............................................................
64_Cyanobium_sp._PCC_7001  MPRRTNRPSTAAPLAPTAPRRRPGVVEAQRGPAAAGAEAAEPAPPLRGTRARAAMQASRT

                                                                                  1
52_Cyanobium_sp._PCC_7001  ........................................................MHPL
64_Cyanobium_sp._PCC_7001  PASPAASRPVGRRSSRPGTTARPASATVRPPRGFSRSAPAASTPAAARSLRANRSRLHPL

                                  10        20        30        40        50        60
52_Cyanobium_sp._PCC_7001  TDASANDALHAYDTAVKLAFDRIVPVLKRLSALQHEDDFVGRAQAIALEELGFPLPEPIL
64_Cyanobium_sp._PCC_7001  TDASANDALHAYDTAVKLAFDRIVPVLKRLSALQHEDDFVGRAQAIALEELGFPLPEPIL

                                  70        80        90       100       110       120
52_Cyanobium_sp._PCC_7001  DTAWVSQLDMRTLYAWCVFETYEQTSEAFFRDDPLQGQPGSPSAEAFDRFLLDCGFHLLD
64_Cyanobium_sp._PCC_7001  DTAWVSQLDMRTLYAWCVFETYEQTSEAFFRDDPLQGQPGSPSAEAFDRFLLDCGFHLLD

                                 130       140       150       160       170       180
52_Cyanobium_sp._PCC_7001  ITPCADGRLAHAIGFGLRLPFSSVRRRPHAGALFDVENTVNRWVKTEHRRYREAQPNPAH
64_Cyanobium_sp._PCC_7001  ITPCADGRLAHAIGFGLRLPFSSVRRRPHAGALFDVENTVNRWVKTEHRRYREAQPNPAH

                                 190       200       210       220       230       240
52_Cyanobium_sp._PCC_7001  ADTRYLKVALYHFSSLDPQHEGCAAHGSDDALAASCGLSRLKDFQQAVENSFCCGASVDL
64_Cyanobium_sp._PCC_7001  ADTRYLKVALYHFSSLDPQHEGCAAHGSDDALAASCGLSRLKDFQQAVENSFCCGASVDL

                                 250       260       270       280       290       300
52_Cyanobium_sp._PCC_7001  LLMGIDTDTDAIRVHVPGMDGSTRLDRWLDARDVYDATLGLPPDQARQRVSALVQEAAAS
64_Cyanobium_sp._PCC_7001  LLMGIDTDTDAIRVHVPGMDGSTRLDRWLDARDVYDATLGLPPDQARQRVSALVQEAAAS

                                 310       320       330       340       350       360
52_Cyanobium_sp._PCC_7001  VPDPGMVTLVARLFEHNISQIDYVRQFHGGAYDDAGHAERFIGVGIGFKEIHLRNLTYFA
64_Cyanobium_sp._PCC_7001  VPDPGMVTLVARLFEHNISQIDYVRQFHGGAYDDAGHAERFIGVGIGFKEIHLRNLTYFA

                                 370       380       390       400       410       420
52_Cyanobium_sp._PCC_7001  YMDTVEEGAADLDVGVKIFKGLNVSRGLPVPVVVRFDYHGQVPGARDRAVRHCQRVQTAI
64_Cyanobium_sp._PCC_7001  YMDTVEEGAADLDVGVKIFKGLNVSRGLPVPVVVRFDYHGQVPGARDRAVRHCQRVQTAI

                                 430       440       450       460
52_Cyanobium_sp._PCC_7001  ESRYPELFQQGLLHALLTVRDQDRHTPAEAVGSTIVFATGGGH
64_Cyanobium_sp._PCC_7001  ESRYPELFQQGLLHALLTVRDQDRHTPAEAVGSTIVFATGGGH
```

**Figure S1**. The full length CyCsoSCA protein contains an extended disordered N-terminal tail for the first 116 residues. The alignment pictured above details the full length CyCsoSCA protein sequence (64 Cyanobium sp. PCC 7001) aligned with the truncated version expressed in this work (52 Cyanobium sp. PCC 7001).

**Table S1**. Data collection and refinement details for the *Cy*CsoSCA structure presented in this work.

| **PDB ID** | **8THM** |
|---|---|
| **Data collection** | |
| Space group | P 21 21 21 |
| Cell dimensions | |
| a, b, c (Å) | 104.314, 181.848, 190.057 |
| a, b, g (°) | 90, 90, 90 |
| Resolution (Å) | 131.392-2.300 |
| $R_{merge}$ | |
| Within I+/I- (overall, inner, outer) | 0.250, 0.052, 4.833 |
| All I+ and I- (overall, inner, outer) | 0.260, 0.0.54, 5.093 |
| $R_{pim}$ | |
| Within I+/I- (overall, inner, outer) | 0.107, 0.022, 2.031 |
| All I+ and I- (overall, inner, outer) | 0.077, 0.017,1.468 |
| I/σI | 8.1, 32.3, 0.7 |
| $CC_{1/2}$ | 0.997, 0.999, 0.297 |
| Completeness (%) | 100.0, 98.3, 100.0 |
| Redundancy (multiplicity) | 12.3,10.4,12.7 |
| **Refinement** | |
| Resolution (Å) | 47.514-2.3 (2.382-2.300) |
| No. reflections | 160413 (15835) |
| $R_{work}/R_{free}$ | 0.1827/0.2392 |
| No. atoms | 23052 |
| Protein | 21974 |
| Ligand/ion | 282 |
| Water | 796 |
| B-factors (overall) | Mean(55.7), max(129.2), min(31.1) |
| Protein | Mean(55.7) max(129.2), min(31.1) |
| Ligand/ion | Mean(61.3), max(95.9), min(40.2) |
| Water | Mean(55.2), max(95.3), min(36.3) |
| R.m.s. deviations | |
| Bond lengths (Å) | 0.0090 |
| Bond angles (Å) | 1.24 |

**Table S2.** Details of the RuBP binding site for each chain of the *Cy*CsoSCA structure presented in this work.

| Chain | $PO_4$ head 1 | C-chain | $PO_4$ head 2 |
|---|---|---|---|
| **A** | R266 (sidechain and backbone), K469, water | 4xwaters, K469 | Water, R560, D517 |
| **B** | R266(sidechain and backbone), water | 4xwaters, R265, D517 | R265 |
| **C** | K469 | 3xwaters, K469 | R560, water |
| **D** | R266, H356, K469 (backbone) | K469, R149 | R560, K469, D517 |
| **E** | R266, water | R265, K469 | R560, D517, K469 |
| **F** | 2xwaters, K469 | R149, 2xwaters, R560 | R560, 2xwaters |

**Table S3**. Residues involved in binding sulfate ions within each chain of the *Cy*CsoSCA structure presented here.

| Chain | $SO_4$ no.1 | $SO_4$ no. 2 | $SO_4$ no. 3 |
|---|---|---|---|
| **A** | R265, 2xwaters, N278 | 2xwater, R267, N278, K307, H269 | R266, water, 2 bbone interactions |
| **B** | R265, 3xwater, R282 | 2xwater, R267, N278, H269, R282 | |
| **C** | R265, K285, R282, 2xwaters, R151 | 2xwaters, H269, N278, R267, R282 | |
| **D** | R265, R282, K285 | H269, R282, N278, R267, water | R266, F468 (backbone), I471 (backbone), water |
| **E** | R265, K285, R282 | R282, N278, R267, H269, water | 2xwater, R266, F468 (backbone) |
| **F** | K285, 2xwaters, R265, R282 | R282, 2xwaters, H269, R267, N278 | R266, 2xwaters, F468 (backbone), I471 (backbone) |

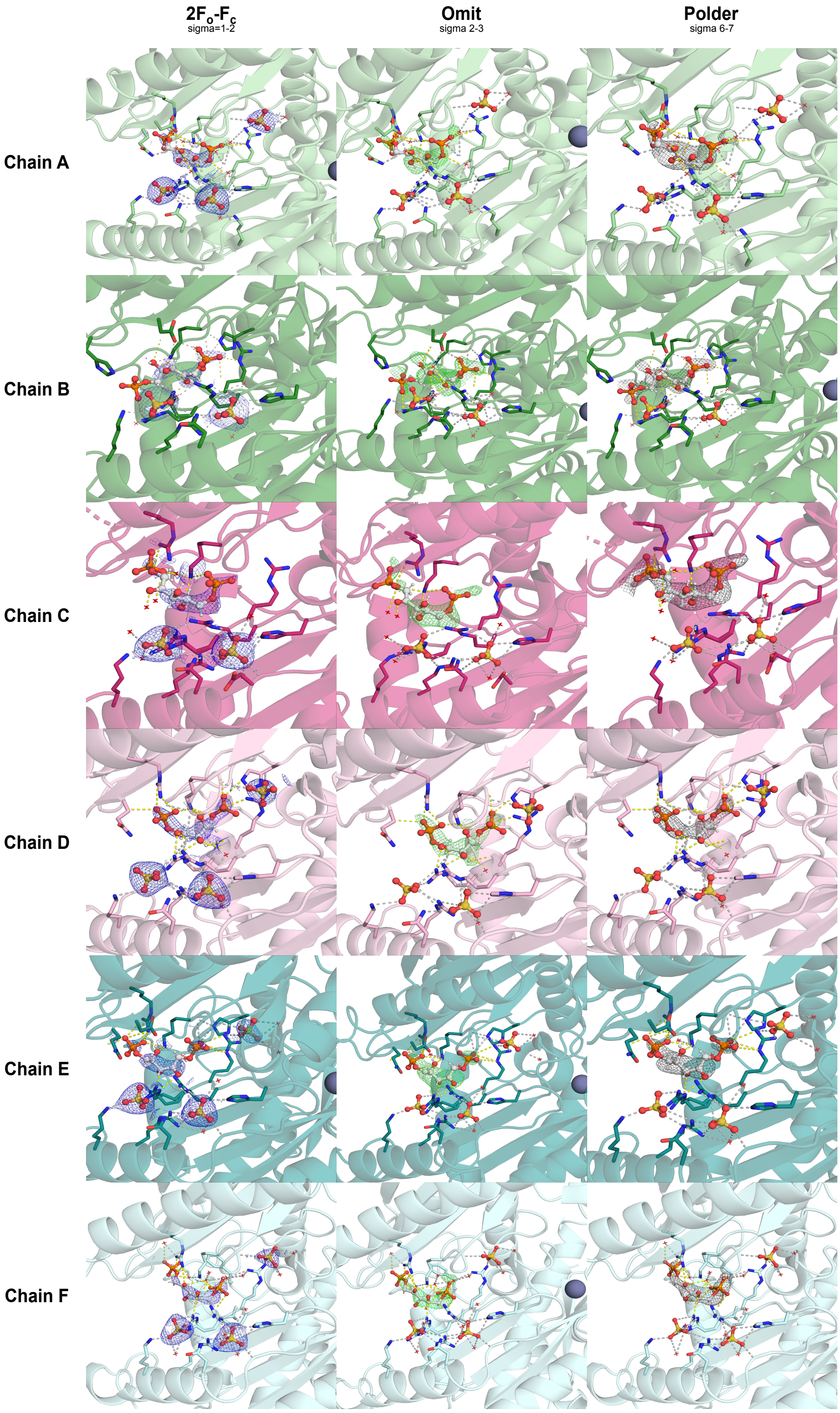

**Figure S2** Density maps of RuBP in each monomer in the *Cy*CsoSCA structure characterised in the main text. At least two sulfate ions are observed at this site in all monomers from the crystallisation solvent (Supp. Figure 1). Sulfate ions have been observed bound to β-CA oligomeric interfaces, though rarely at pockets buried so deep as observed here[1].

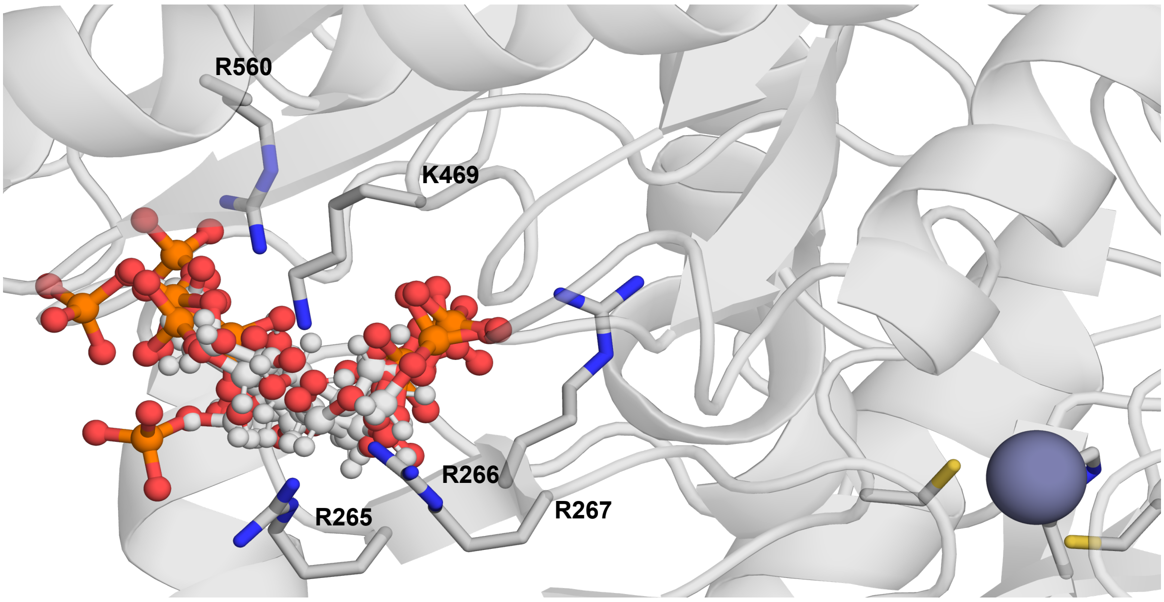

**Figure S3.** RuBP molecules in each monomer overlayed. The ligand is shown in ball and stick representation and coloured by element. Key sidechains are shown in stick representation and annotated. For perspective, the overlayed ligands have been situated in chain D (cartoon representation) with the zinc active site shown as a grey sphere.

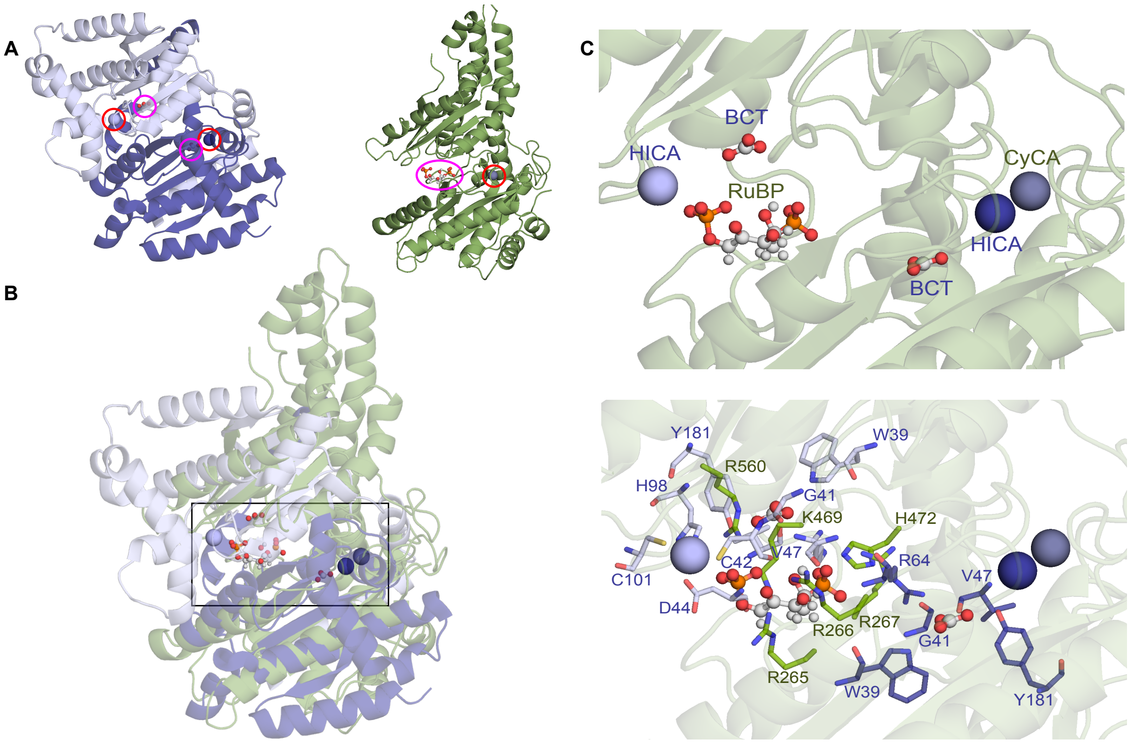

**Figure S4.** The RuBP binding site is distinct from previously identified allosteric bicarbonate site documented in Type II β-CAs, typified by the Haemophilus influenzae (HICA; PDB:2A8D)[1–5]. **A** The HICA dimer shown with monomers in different hues of purple and the *Cy*CsoSCA monomer (chain D shown here) in green. The allosteric ligands are indicated by a magenta circle and the active zinc residues by a red circle in each structure. Type II β-CAs exhibit a pH-dependent activity profile symptomatic of an allosteric bicarbonate inhibition mechanism. Broadly, bicarbonate binding causes a conformational shift that disrupts the conserved Asp-Arg active site dyad, causing the Asp to coordinate the zinc ion and displace the catalytically essential water molecule[1]. The allosteric RuBP (*Cy*CsoSCA) and bicarbonate ions (HICA) are shown in ball and stick representation. Zinc ions are shown as spheres. B A structural alignment of the HICA dimer and the *Cy*CsoSCA monomer is shown with a box highlighting the zinc active sites and ligand binding pockets. **C** A zoomed in box of the overlayed zinc ions and allosteric ligands. Zinc ions corresponding to HICA or *Cy*CsoSCA (*Cy*CA) are shown as spheres. The RuBP ligand associated with *Cy*CsoSCA and bicarbonate ions (BCT) associated with HICA are shown in ball and stick representation and annotated. In the box below, ligand binding residues are shown in stick representation, purple residues correspond to sites in the HICA structure and green residues those in the *Cy*CsoSCA structure. The catalytic zinc binding residues C42, D44, H98 and C101 are shown the HICA monomer closest to the *Cy*CsoSCA RuBP binding site. Though in a similar region of the protein, the two sites employ different residues and the RuBP sits much further from the active site relative to the allosteric bicarbonate site. These differences are largely due to the lack of active site pairing in the CsoSCA pseudo-dimer. While the fundamental unit of β-CAs is typically a dimer with symmetric active site pairing (or a pseudo-dimer in which the two monomers have fused but the active sites are each maintained), the CsoSCA clade has lost this defining feature[3]. Here, the C-terminal domain (CTD) does contain key structural motifs reminiscent of the Catalytic domain but has diverged so significantly that the active site is no longer intact.

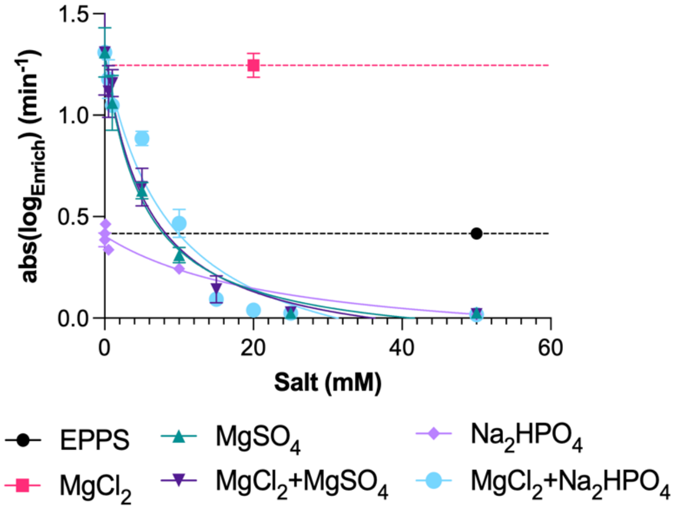

**Figure S5.** Given the overlaying positions of RuBP and sulfate ions, we hypothesised sulfate may compete with RuBP at this site, explaining low ligand density despite high concentrations in crystallisation conditions. *Cy*CsoSCA activity in the presence of sulfate and phosphate ions was quantified using the MIMS[8] as detailed in the main text. All assays are run in the presence of 20mM EPPS. Activity in the absence of any salt (20mM EPPS) is shown, highlighting the requirement for $Mg^{2+}$ to achieve maximal activity (50mM $MgCl_2$). Activity levels at this basal condition are indicated as single points/dashed horizontal lines. CA activity detected at increasing concentrations of key salts $MgSO_4$ and $Na_2HPO_4$ as shown in the figure legend are plotted. These results show *Cy*CsoSCA is significantly inhibited at 50mM $MgSO_4$ relative to standard assay conditions (20mM $MgCl_2$). Phosphate ions, a more biologically relevant ion with analogous size and charge, had a comparable effect to sulfate. *Cy*CsoSCA activity also dropped in the absence of Magnesium salts, perhaps functioning to stabilise the ligand in solution. It is unclear whether phosphate would ever reach millimolar concentrations within the carboxysome *in vivo*. All data points are the mean of three replicates, error bars denote standard error of the mean.

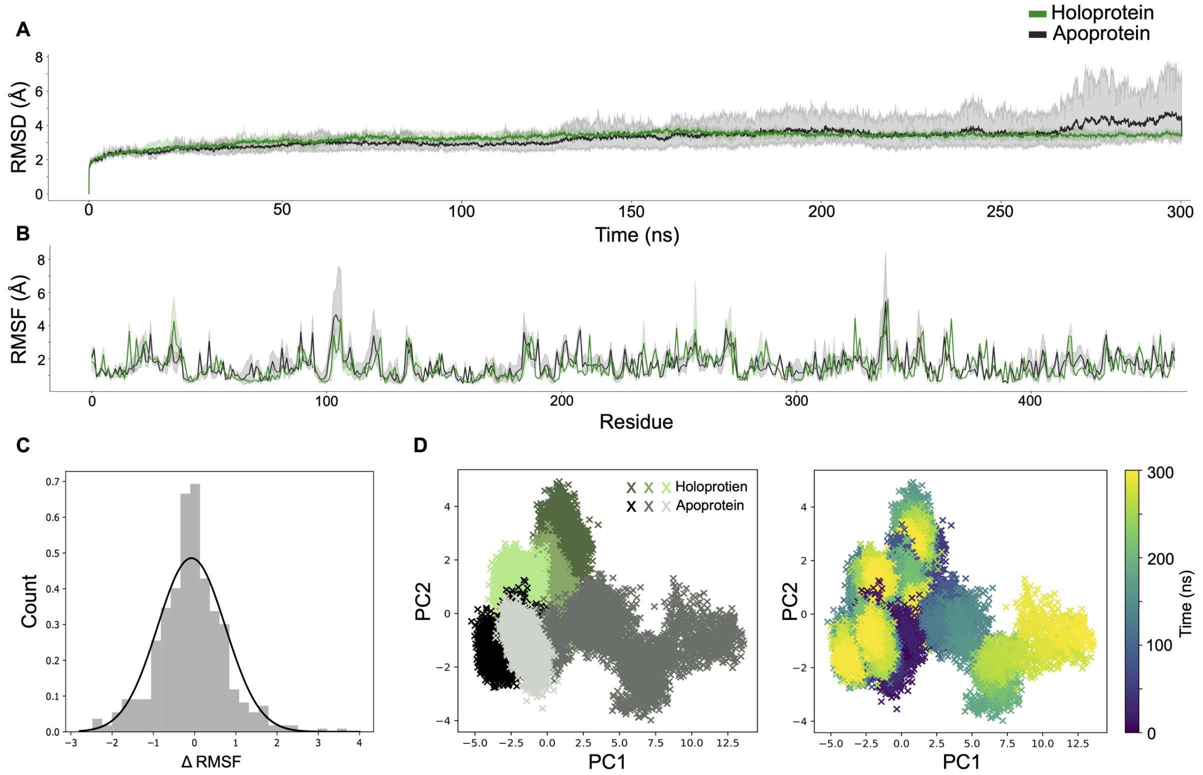

**Figure S6**. Trajectory analysis of molecular dynamic simulations of CyCsoSCA Chain D with (holoenzyme) and without (apoenzyme) RuBP present. **A** Cα RMSD (Å) variation across trajectory for each 300ns replicate relative to frame 1. Three replicates were conducted for each condition with frame recording intervals of 15ns, error indicates standard deviation. Both trajectories equilibrated, with a few fluctuations evident in the apoprotein trajectories particularly after 250ns. **B** Cα RMSF (Å) of each residue across the trajectory highlighting marginal site-by-site differences. **C** Histogram recording the difference between RMSF of analogous residues. Average RMSF (Å) was calculated for each residue and values from the apoenzyme trajectory were subtracted from the average RMSF value of the corresponding residue in the holoenzyme trajectory. Thus, a negative value is indicative of a residue with a greater RMSF in the apoenzyme trajectory and vice versa. These values have then been plotted onto the structure. **D** Principal component analysis of cartesian coordinates across all replicates coloured by either replicate (different shades of green are indicative of holoenzyme replicates, shades of grey indicate apoenzyme replicates) or time within replicate trajectories (scree test: 0.44, 0.12). This shows that while all seem to start in relatively similar conformations, holoprotein trajectories vary more along PC2 over time.

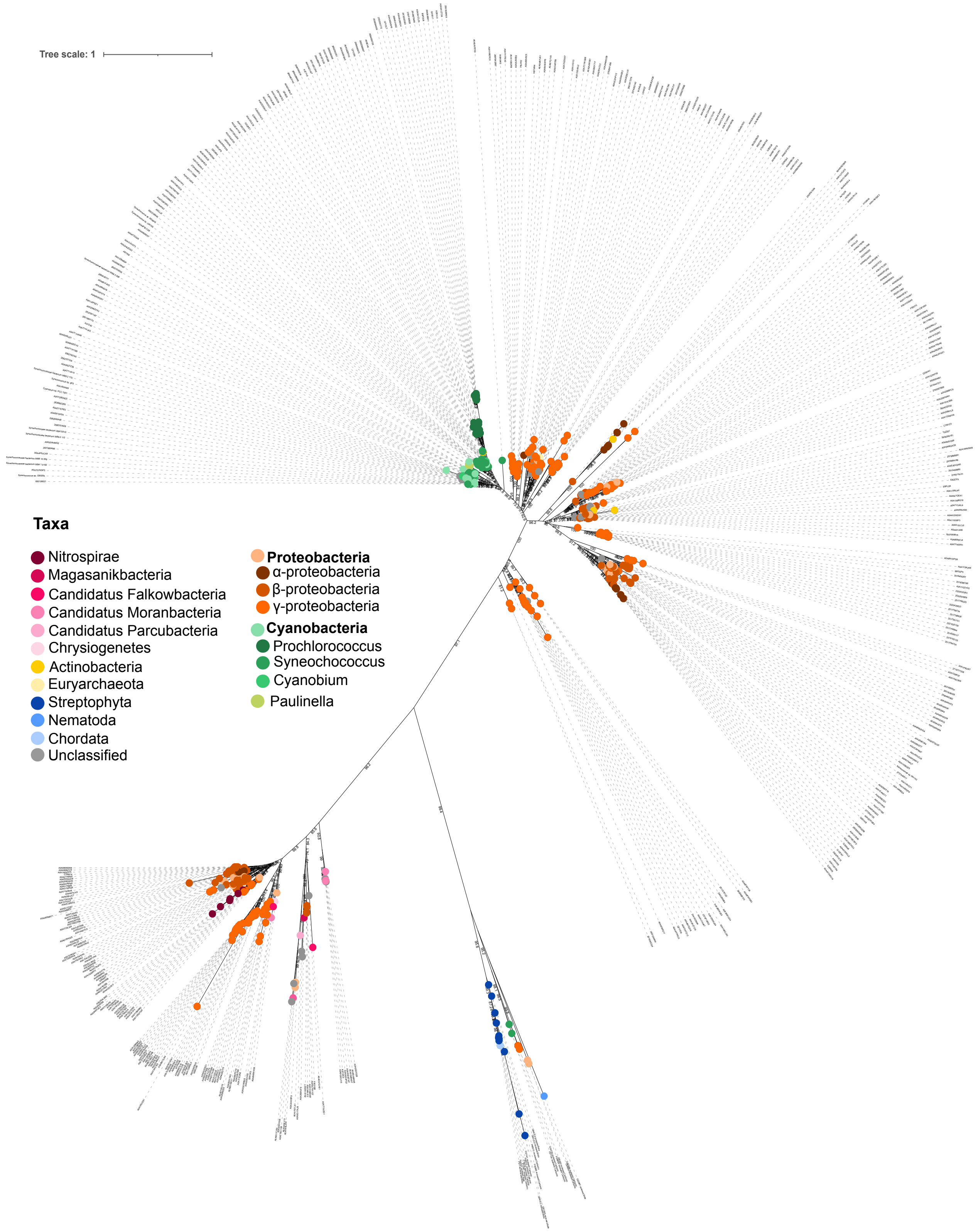

**Figure S7.** An unrooted phylogeny of the CsoSCA protein family inferred by maximum likelihood generated through IQ-Tree using standard parameters. Bootstrap values >70 generated using the ultrafast bootstrap approximation form 1000 replicates[9]. Extant nodes coloured according to taxa as per key in the legend.

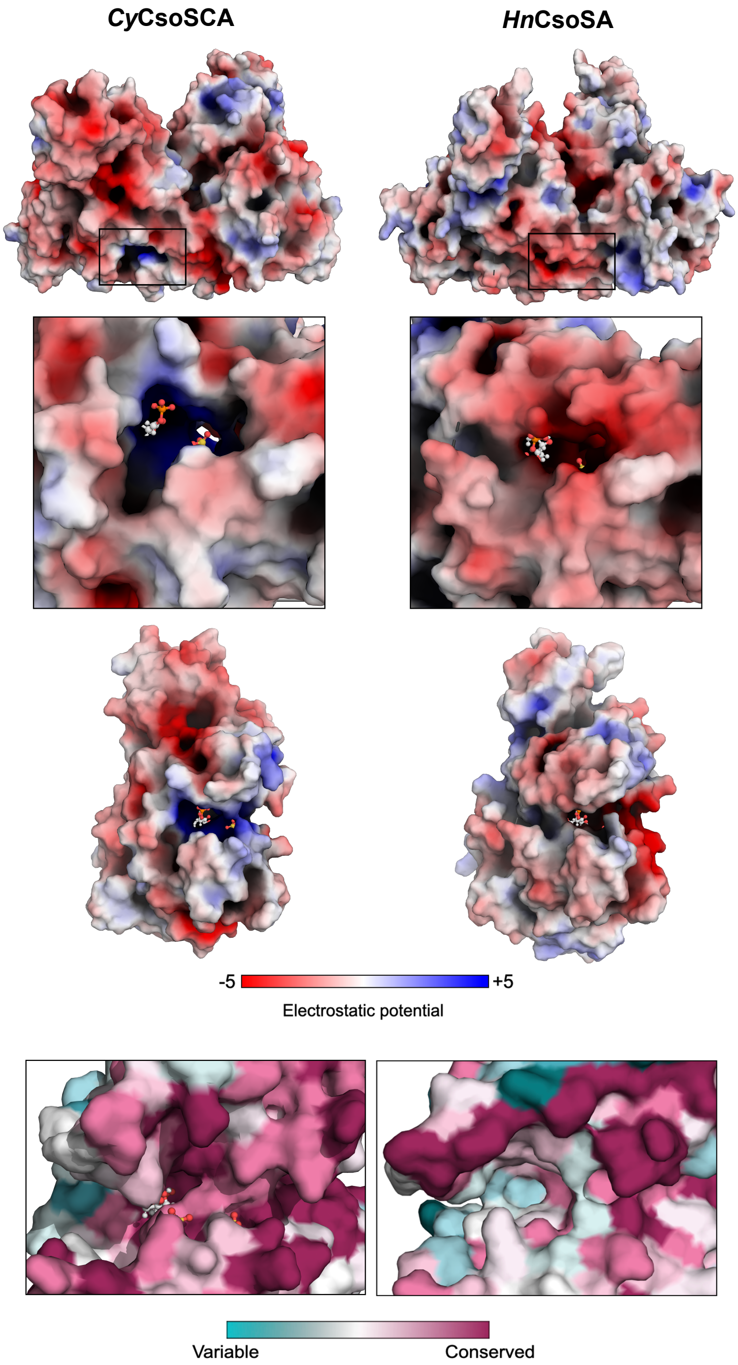

**Figure S8.** The RuBP pocket is conserved and positively charge in cyanobacterial CsoSCA variants. *Cy*CsoSCA and *Hn*CsoSCA structures coloured by electrostatic charge and residue conservation as per legends in the figure. From top to bottom, the dimer of each variant is shown, followed by a zoomed in box of the RuBP binding site, and then a view of the ligand pocket in the monomer. Given RuBP primarily exists in a negatively charged state[18], binding site charge was of key interest. In *Cy*CsoSCA, this site is highly positively charged in *Cy*CsoSCA while the comparative region in the constitutively active *Hn*CsoSCA has a negative electrostatic potential. The final panel displays a zoomed in box of the ligand pocket in each variant coloured by conservation. RuBP has been aligned onto the *Hn*CsoSCA for perspective. The multiple sequence alignment (MSA) of the curated CsoSCA dataset, comprising 134 cyanobacterial species and 337 other taxa, was used to assess the conservation of key features associated with RuBP binding across the protein family. Separate alignments of each taxonomic group were constructed and submitted to the ConSurf webserver for image generation[10]. The resulting conservation values were plotted onto the *Cy*CsoSCA structure and the *Hn*CsoSCA structure for cyanobacterial and other taxa alignments respectively. The positive charge lining the RuBP pocket in *Cy*CsoSCA appears to be a highly conserved feature among analysed cyanobacterial homologues. Comparatively, the conservation of the equivalent region in other taxa to the *Hn*CsoSCA structure, demonstrates the strong negative charge is not a conserved feature and that this region is quite variable across other taxa surveyed. This data is consistent with the positive charge of the region, and likely the capacity for RuBP binding, being conserved across cyanobacterial CsoSCAs but absent or un-conserved in other taxa.

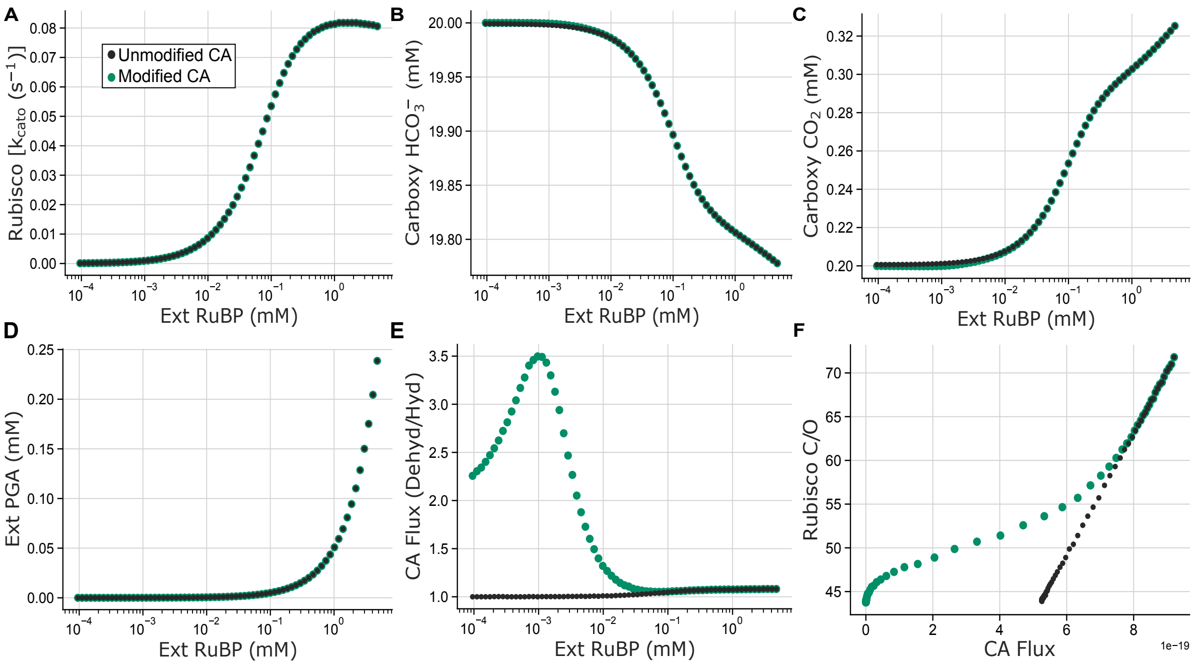

**Figure S9.** The model presented in Long et al. (2021) simulating carboxysome function was adapted to emulate a *Cyanobium* α-carboxysome with either an RuBP-dependent CA (Modified CA, green dots) or a constitutively active CA (Unmodified CA, black dots). **A** Rubisco oxygenation activity per second (Rubisco (kcato) is equivalent between the two systems across modelled cellular RuBP concentrations (Ext RuBP (mM)), corresponding to the equivalent observed carboxylation rates (Figure 5). **B** The concentration of $HCO_3^-$ and **C** $CO_2$ within the carboxysome as a function of modelled cellular RuBP concentrations (Ext RuBP (mM)). **D** Correspondingly, the concentration of modelled cellular PGA (Ext PGA (mM)) as a function of modelled cellular RuBP concentrations (Ext RuBP (mM)) demonstrates no difference in PGA production between the two systems. **E** While the carboxysomal CA is capable of both the forward and reverse $CO_2$ hydration/$HCO_3^-$ dehydration reactions, to supply the carboxysome-encased Rubisco with $CO_2$ requires the dehydration reaction to predominate. The ratio of the fluxes recorded for the CA dehydration to hydration reaction (CA flux (Dehyd/Hyd)) is plotted here across a modelled cellular RuBP gradient (Ext RuBP (mM)). This demonstrates notable differences in CA flux and at low RuBP conditions in the two systems. As Ci appears consistent between each system, this likely manifests as changes in the carboxysomal proton concentration at these RuBP levels. Indeed, this is consistent with the observed changes in pH presented in Figure 5C. **F** The ratio of Rubisco carboxylation rates to oxygenation rates (Rubisco C/O) as CA activity (CA flux, indicative of CA activity levels converting $HCO_3^-$ to $CO_2$) increases with cellular RuBP concentrations in the modified and unmodified systems.

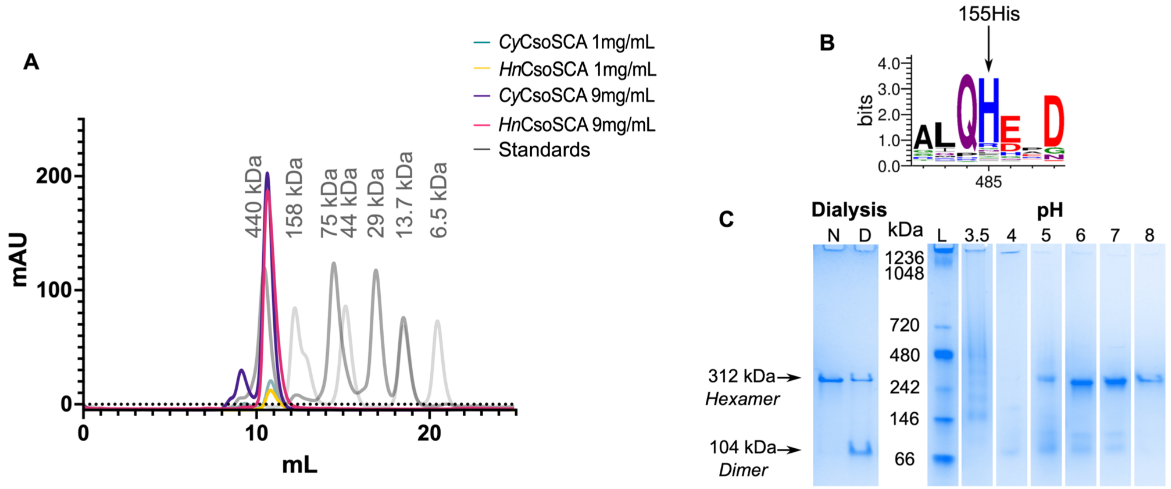

**Figure S10.** A homohexameric trimer of dimers is the prevailing, biologically relevant form of CsoSCA. **A** Size exclusion chromatography of *Cy*CsoSCA and *Hn*CsoSCA shows each isoform elutes at approximately the volume expected for the hexamer observed in the *Cy*CsoSCA crystal structure (approx. 312 kDa) independent of protein concentration. An additional peak in the *Cy*CsoSCA samples at both concentrations is also seen at slightly higher molecular weights suggesting, while the hexamer is the predominant form, a spectrum of oligomeric states may be present in solution. **B** A sequence logo with the key His155 residue required by all monomers to coordinate the structural zinc ion in an octahedral $His_3(H_2O)_3$ coordination sphere (Figure 1 main text). The key zinc coordination residue appears conserved across all analysed CsoSCA sequences. **C** To assess the role of the zinc ion in coordinating this observed hexameric state, we perturbed zinc ligand binding and observed the effect on oligomerisation. Native *Cy*CsoSCA (N) and *Cy*CsoSCA dialysed against 2mM 1,10-phenanthroline, a string chelating agent, for 24 hours before being run on a Native PAGE to compare oligomeric states. This results in the breakdown of the high molecular weight band corresponding to the hexameric state, in favour of a smaller band approximately the size expected for the dimer. Additionally, was *Cy*CsoSCA incubated across a pH gradient and subsequently assessed by Native PAGE to capture pH dependent perturbations in oligomeric state. as solution pH approaches the pKa of Histidine a corresponding oligomeric shift is observed. At pH 3.5 smearing is indicative of protein denaturation and at pH 4 a faint band is visible between 146 kDa and 66 kDa consistent with the dimeric *Cy*CsoSCA (104 kDa). In samples incubated at pH 5, the dimeric band increases in intensity and a second band is evident between 242 kDa and 480 kDa consistent with the hexameric *Cy*CsoSCA (312 kDa). As the pH becomes more basic the hexameric band increases in intensity and the dimeric band fades. This would correspond with the hexamer destabilising as the Histidines coordinating the structural Zinc ions are protonated, and dimers become the prevailing quaternary state in solution. At pH 6 and 7 a second faint band is evident above the dimeric band between the 66kDa and 146kDa markers. The distance between the two bands here is too small to denote a trimer thus it is likely this is some kind of contaminant. This data is consistent with the NTD acting as an oligomerisation domain, binding structural Zincs to facilitate the formation of large hexameric CsoSCA assemblies in solution. The standard is labelled (L) with corresponding weights annotated in kDa.

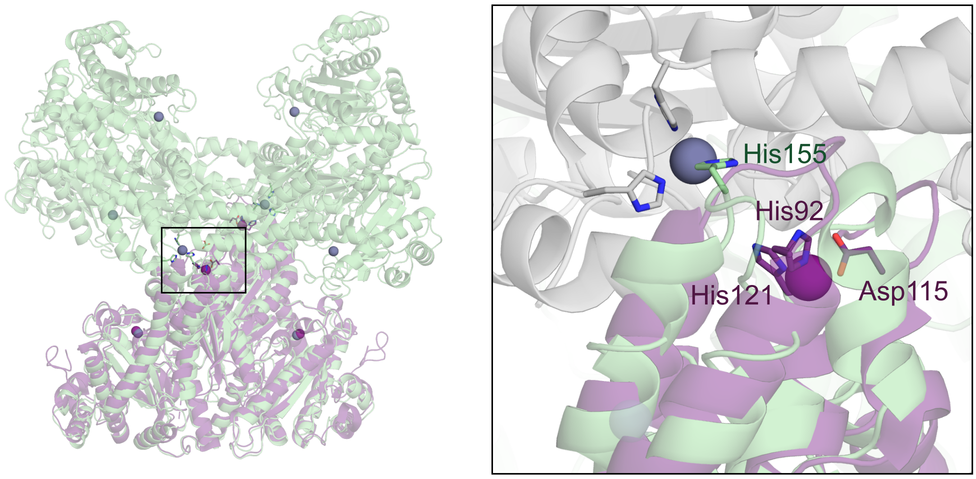

**Figure S11.** The opportune zinc site in the NTD of the crystallised *Hn*CsoSCA previously characterised by Sawaya et al. (2006) arose from an inadvertent mutant and likely precluded the formation of the true hexameric state[21]. The HnCsoSCA dimer (PDB:2FGY) is shown (purple) overlaid with the *Cy*CsoSCA structure solved here (green) with zinc ligands shown as spheres (grey spheres for *Cy*CsoSCA associated Zincs and purple spheres for those associated with HnCsoSCA). A box with a zoomed in view of the structural zinc site is shown with key zinc binding ligands in *Cy*CsoSCA (His155) and *Hn*CsoSCA (His92, Asp115, His121) shown in stick representation. The purple sphere shown here depicts the non-biologically relevant zinc-binding site in the NTD while the grey sphere denotes the additional zinc ion demonstrated to mediate additional contacts necessary for hexamer formation.

**Table S4.** DALI search results for a non-*cso* sequence (UniProt ID A0A2N1ALC1)

| Chain | Z score | RMSD | Lali | No.res | %ID | Description |
|---|---|---|---|---|---|---|
| **2fgy-a** | 31.4 | 2.8 | 340 | 471 | 18 | Carboxysome shell polypeptide |
| **1ddz-a** | 17.5 | 3.4 | 273 | 481 | 14 | Carbonic anhydrase; |
| **3vrk-a** | 10.7 | 2.7 | 128 | 212 | 12 | Carbonyl sulfide hydrolase; |
| **3vqj-a** | 10.6 | 2.7 | 128 | 213 | 12 | Carbonyl sulfide hydrolase; |
| **6gwu-b** | 10.6 | 3.4 | 141 | 205 | 11 | Carbonic anhydrase; |
| **2a5v-c** | 10.4 | 2.7 | 131 | 201 | 6 | Carbonic anhydrase (carbonate dehydratase)(carbo) |

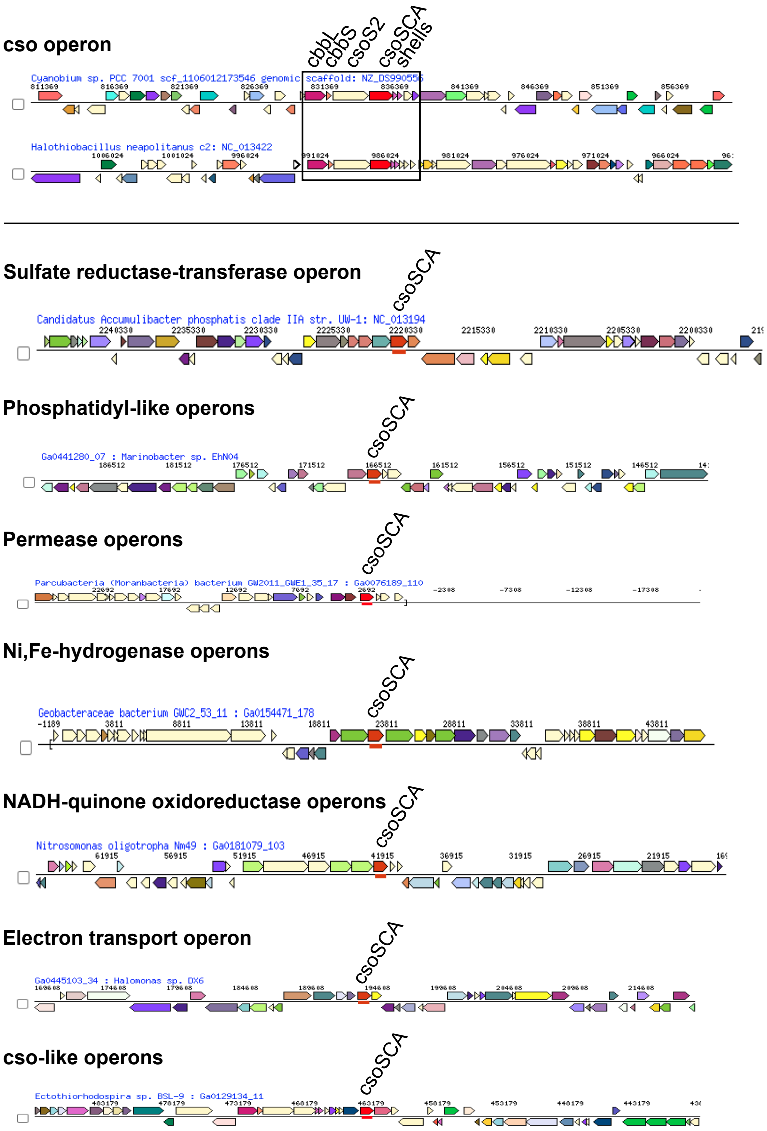

**Figure S11.** Example of operon types observed in CsoSCA dataset. For each sequence in the 'non-cso' cluster the genomic context was pulled using the JGI IMG database and manually assessed. Canonical *Hn*CsoSCA and *Cy*CsoSCA demonstrate typical cso operon structure. The colours of each gene are automatically generated on JGI IMG based on different COG annotations. The majority of 'non-cso' sequences could be classed into the 7 listed operon types (arbitrarily named based on key functional annotations). Full details of operons in attached files.

**Table S5.** Primers used in this work to sequence and generate *Cy*CsoSCA mutants.

| **Target gene** | **Purpose** | **Destination Vector** | **Primer** | **Sequence** |
|---|---|---|---|---|
| Cyanobium csoSCA | epPCR | pET16 | csoS3_epPCR_F_BsaI | TAGGTCTCTCATATGCCGCGTCGTACC |
| | | | csoS3_epPCR_R_BsaI | TAGGTCTCTTCGATTAGTGACCGCCGCC |
| | Sequencing epPCR mutants | | pET16_Seq_F | ATCTCTATATCTCTATTTGACGGCT |
| | | | pET16_Seq_R | CTAGGGCAAACATATAAACGCA |
| | | | CyCsoS3_sR1 | TTACGCCAACATCCAGGTCTGC |
| | Creating H519Q CyCsoSCA mutant | pHue | Gibson_CA52_H519Q_F | CGTTTTGACTATCAGGGTCAGGTACCAGG |
| | | | Gibson_CA52_H519Q_R | CCTGGTACCTGACCCTGATAGTCAAAACG |
| | Creating T477A CyCsoSCA mutant | | Gibson_CA52_T477A_F | GCGTAATCTGGCATACTTCGCTTACATGG |
| | | | Gibson_CA52_T477A_R | CCATGTAAGCGAAGTATGCCAGATTACGC |
| | Creating N278D CyCsoSCA mutant | | Gibson_CA52_N278D_F | TGACGTTGAAAAAACCGTGAACCGTTGG |
| | | | Gibson_CA52_N278D_R | CCAACGGTTCACGGTTTTTTCAACGTCA |
